# Identifying flow modules in ecological networks using Infomap

**DOI:** 10.1101/2020.04.14.040519

**Authors:** Carmel Farage, Daniel Edler, Anna Eklöf, Martin Rosvall, Shai Pilosof

## Abstract

1. Analysing how species interact in modules is a fundamental problem in network ecology. Theory shows that a modular network structure can reveal underlying dynamic ecological and evolutionary processes, influence dynamics that operate on the network and affect the stability of the ecological system.
2. Although many ecological networks describe flows, such as biomass flows in food webs or disease transmission, most modularity analyses have ignored network flows, which can hinder our understanding of the interplay between structure and dynamics.
3. Here we present Infomap, an established method based on network flows to the field of ecological networks. Infomap is a flexible tool that can identify modules in virtually any type of ecological network and is particularly useful for directed, weighted and multilayer networks. We illustrate how Infomap works on all these network types. We also provide a fully documented repository with additional ecological examples. Finally, to help researchers analyse their networks with Infomap, we introduce the open source R package infomapecology.
4. Analysing flow-based modularity is useful across ecology and transcends to other biological and non-biological disciplines. A dynamic approach for detecting modular structure has strong potential to provide new insights into the organisation of ecological networks.

## Introduction

Understanding the interplay between the structure and dynamics of complex ecological systems is at the heart of network ecology. Partitioning a network into modules composed of nodes more tightly connected to each other than to other nodes is a leading example. While there is little empirical evidence for the physical existence of modules, the theoretical understanding of modularity is expansive. It has been shown that a modular structure can make ecological communities locally stable (Grilli *et al.*, 2016), increase species persistence (Stouffer & Bascompte, 2011), serve as a signature for evolutionary processes (*Pilosof et al.*, 2019) and slow down the spread of perturbations (see Gilarranz *et al.* 2017 for experimental evidence).

There are three main ways to detect modules in networks (Rosvall *et al.*, 2018): (i) By maximising the internal density of links within groups of nodes (Newman & Girvan, 2004; Olesen et al., 2007; Thébault, 2013); (ii) by identifying structurally equivalent groups in which nodes connect to others with equal probability, typically studied using stochastic-block models (Holland et al., 1983), known as the ‘group model’ in ecology (Allesina & Pascual, 2009); and (iii) by optimally describing modular flows on networks (Rosvall & Bergstrom, 2008; Rosvall et al., 2010) (Table SI 1). These approaches have been developed for different purposes, with different mathematical functions and algorithms to detect an ‘optimal’ partition of a network. Therefore, there is no single ‘true’ network partition (Peel et al., 2017). Instead, the method applied should match the question (Ghasemian et al., 2019; Rosvall et al., 2018). For example, many ecological systems describe flows on networks, including biomass flow in food webs (Baird & Ulanowicz, 1989), movement of individuals between patches (Hanski & Gilpin, 1991) and gene flow among individuals and populations (Fletcher, Jr et al., 2013). In such cases, understanding how network flows organise in modules can be more relevant to the system at hand than maximising internal density.

To date, maximising variants of Newman-Girvan’s combinatorial modularity score *Q* is the dominant approach in ecology (reviewed in Thébault (2013)). While this method undoubtedly has provided many insights, it is not designed to capture network flows. Also, modularity maximisation methods for various applications are scattered in different software implementations. For example, the R package biparite (Dormann et al., 2009) has an implementation for modularity maximisation in biparite weighted and unweighed networks, while Netcarto (Guimerà & Nunes Amaral, 2005) is an implementation for unipartite, undirected networks. To fill these conceptual and technical gaps, we present an established method for detecting flow-based modules called Infomap.

Infomap has several advantages for ecological research. First, it can be applied to many types of networks, including directed/undirected, weighted/unweighted, unipartite/bipartite, and multilayer networks. Second, it is computationally effective, supporting studies of large networks or comparing observed networks with many randomised networks. Third, it can incorporate node attributes by explicitly considering information such as taxonomy in the partitioning to modules. Fourth, it can detect hierarchical structures of modules within modules. Finally, Infomap has online documentation and an active development team that has made it user-friendly and flexible. These advantages make Infomap a highly accessible tool that can be applied to virtually any kind of ecological system. Moreover, Infomap has been thoroughly described mathematically and computationally (Rosvall & Bergstrom, 2008; Rosvall et al., 2010; Rosvall & Bergstrom, 2011; Rosvall et al., 2014), and has already been benchmarked against other methods (Lancichinetti & Fortunato, 2009; Aldecoa & Marín, 2013), providing a sound theoretical and applied understanding of the method.

Despite these advantages, Infomap has only been used in a handful of ecological studies (Pilosof et al., 2019; Bernardo-Madrid et al., 2019; Pilosof et al., 2020). Therefore, our goal here is twofold: (i) introduce Infomap to ecologists with guidelines on how to apply it to particular problems; and (ii) help users analyse their networks with the dedicated R package infomapecology we have developed – a one-stop-shop that also integrates with other R packages commonly used by ecologists such as bipartite and igraph.

## Infomap and the map equation objective function

### General approach to network partitioning

To understand how Infomap works, it is helpful first to understand the general approach for modularity analysis (SI 2). A particular assignment of the nodes into modules is called a network partition. As even small networks can have an enormous number of possible partitions, search algorithms measure the quality of a given partition with an objective function. The algorithms then make a small change in the partition, such as moving a node from one module to another, and test if the value of the objective function improves. Modularity analysis algorithms differ in the search algorithms and objective functions they apply.

Infomap optimises the objective function known as the map equation using a modified and extended Louvain search algorithm (Blondel et al., 2008). Specifically, the algorithm finds the partition that best compresses a description of flows on the network. The network flows are modelled by a random walker or observed empirical flows if available (SI 3). The random walker moves across nodes in a way that depends on the direction and weight of the links, and tends to stay longer in dense areas that then represent modules. For a given partition of the network, there is an associated information cost, measured in bits, for describing the movements of the random walker. The map equation converts the flow rates within and between the modules to an information-theoretic modular description measure of the random walker’s movements on the network. Minimising the map equation over possible network partitions corresponds to detecting the most possible modular structure in the dynamics on the network.

### The map equation: linking structure and information

To calculate the map equation, Infomap uses node and link rates, which are calculated based on link direction and weights. For example, in the schematic network in Fig. 1a, there are 14 directed links of weight of 1, resulting in total incoming link weight of 14. Therefore, each directed link carries flows of link visit rate 1/14. These can also be viewed as 7 undirected links (flow equals link weights in undirected networks). Nodes with two incoming links have a node visit rate of 2/14, and nodes with three links have a node visit rate of 3/14. These rates are included in a so-called ‘module codebook’. In the one-module solution, all the nodes belong to a single module and, therefore, to a single module codebook (Fig. 1c). In the two-module solution (Fig. 1b), there are two module codebooks (Fig. 1d). To describe a random walk in the latter case, it is also necessary to consider the rates of entering and exiting each module using the module entry rate and the module exit rate, respectively (which are equal for undirected networks). Module entry rates are encoded in an ‘index codebook’. In the two-module solution, these events are “enter green” and “enter orange”, which both occur at rate 1/14. The rates of exiting modules are encoded within the module codebooks (Fig. 1d).

The map equation uses Shannon’s source coding theorem (Shannon, 1948) to convert the rates encoded in the codebooks to information measured in bits. Specifically, given a network partition M, we can calculate the minimum amount of information needed to describe an average movement length of a random walker. This quantity *L* is the entropy *H* of the events encoded in the codebooks, weighted by the use rate of each codebook (equations in Fig. 1c,d). Summing the terms for the index codebook and the module codebooks, we obtain the map equation (Rosvall & Bergstrom, 2008; Rosvall et al., 2010),

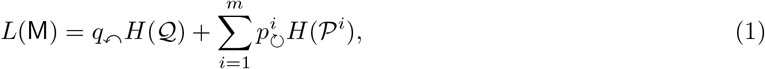

where 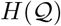 and 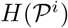 are the entropy values of the index codebook and the codebook of module *i*, respectively. These entropy terms are weighted by the rate at which the codebooks are used. The index codebook is weighted by the rate of entering any module, *q*↶, and each module codebook i is weighted by its within-module flow, 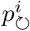, which includes the node-visit rates and the exit rate in module *i*. For the examples in Fig. 1, *L*(M_1_) ≈ 2.56 for the one-module solution and *L*(M_2_) ≈ 2.32 for the two-module solution. The two-module solution requires fewer bits and better captures modular structure in the dynamics on the network.

In practice, Infomap can use either real measured flows, or estimates of flows (SI 3.2). In the latter and more typical case, Infomap derives link and node visit rates using an iterative process akin to the PageRank algorithm (Brin & Page, 1998). First, each node receives an equal amount of flow volume. Then, iteratively until all node visit rates are stable, each node distributes all its flow volume to its neighbours proportionally to the outgoing link weights. We note that PageRank is only used for directed networks because it is superfluous for undirected networks. A comprehensive description on flow models can be found in the Supplementary Information (SI 3.2) and in Rosvall & Bergstrom (2008); Rosvall *et al.* (2010); Bohlin *et al.* (2014); De Domenico *et al.* (2015).

**Figure 1:**
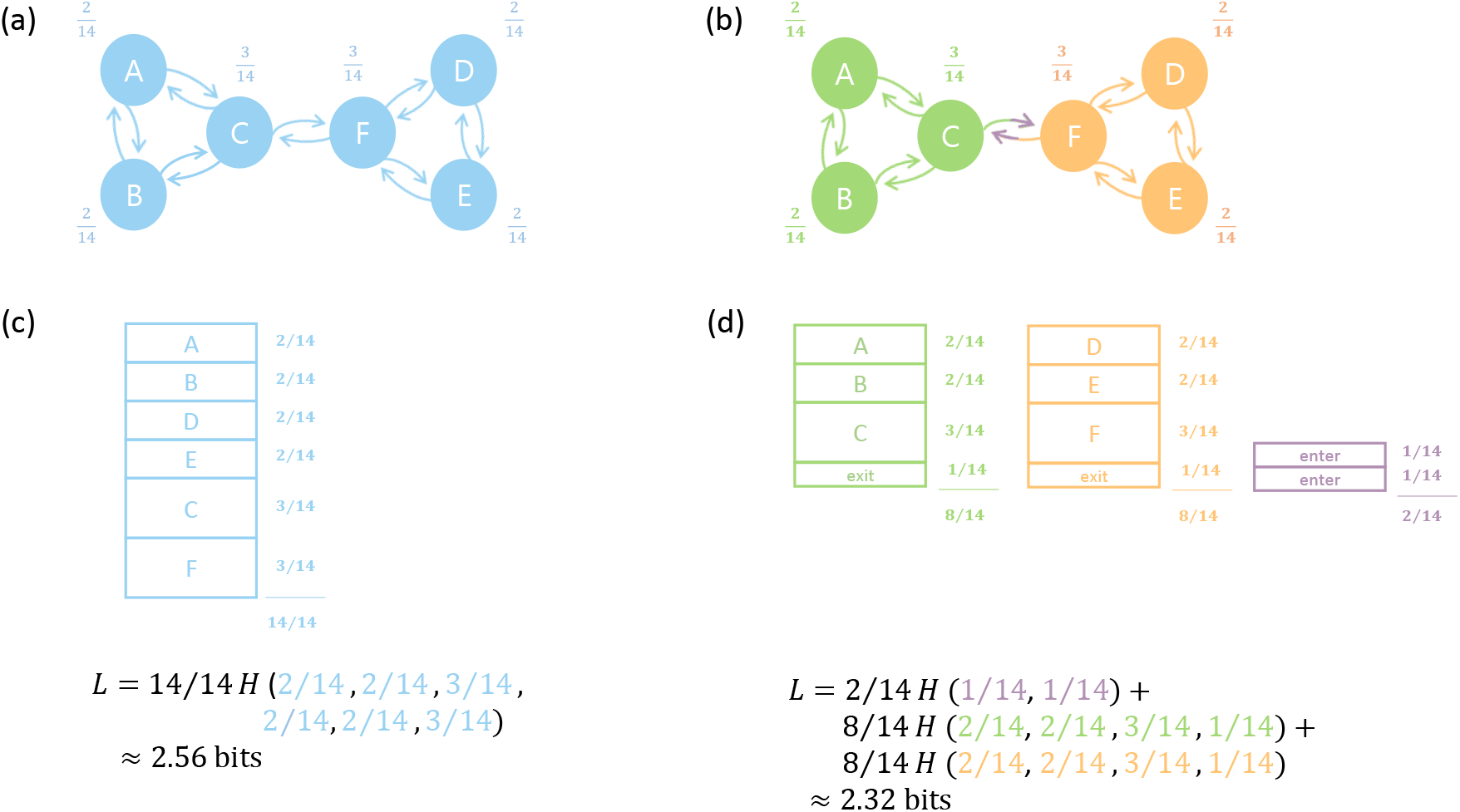
Basics of the map equation. The example is for a schematic network with 14 directional links, in which the total rate of flow is 14. (a) A one-module solution. The node visit rates written besides each node are the number of incoming links. (b) A two-module solution. Each module is represented by a different colour. The purple arrow heads illustrate that these links are considered as module entry links. (c) A single module codebook for a one-module solution. Each block represents a node, with width proportional to the node’s visit rate. (d) Three codebooks for a two-module solution. An index codebook to encode module enter rates (purple codebook), and two module codebooks (green and orange) for node visit rates of each module. The total rate of use of a codebook is the sum of its rates (indicated below the codebook). In (c) and (d), for each solution we calculate *L*, the value of the map equation, which is the entropy of the use rates within each codebook, weighted by the use rates of the codebooks. An expanded explanation that includes the relationship between flow rates and information theory can be found in Fig. S1.

### Extension to multilayer networks

In multilayer networks, nodes representing observable entities such as species are called physical nodes. Realisations of physical nodes in a given layer – for example, in different time points, patches or interaction types – are called state nodes. The random walker moves from state node to state node within and across the layers. However, the encoded position always refers to the physical node (see dynamic visualisation: https://www.mapequation.org/apps/multilayer-network/index.html). This approach provides two advantages. First, it enables a physical node to be assigned to different modules in different layers. From an ecological perspective, this is crucial as a certain species can have different functions in different layers. For example, there is strong spatial and temporal variation in plant-pollinator interactions (Olesen et al., 2008; Trøjelsgaard et al., 2015). Second, it enables to model the coupling between layers without interlayer links. This feature is particularly useful in ecology because interlayer links are often challenging to measure empirically (Hutchinson et al., 2018). If interlayer links are not provided, the random walker ‘relaxes’ to the current physical node in a random layer at a ‘relax rate’ *r*, without recording this movement. By gradually tuning the relax rate, it is possible to explore the relative contribution of intra- and interlayer links to the structure (Fig. 5 and SI 3.4).

## Implementation, availability and code

Full documentation of Infomap, including tutorials, instructions and visualisation tools is available at https://www.mapequation.org/infomap/. Detailed installation instructions for infomap and infomapecology, detailed descriptions of input/output formats, source code of infomapecology, and the code used to produce the analyses in this paper, are available at https://ecological-complexity-lab.github.io/infomap_ecology_package/. In addition, each function in infomapecology has examples in its description, accessible via R’s help (e.g., ?create_monolayer_object).

### General approach

When using infomapecology, the first step is to convert the input data to an object of class monolayer or multilayer. The monolayer class is an R list with information about the network (e.g., bipartite, directed), a list of nodes and their attributes, and network representations as a matrix, an edge list and an igraph object. With multiple data structures, it is easy to streamline and standardise the workflow with other R packages. As ecological networks are typically relatively small, using multiple data structures have limited computational consequences. If the network is large, it is straightforward to extract only a single data structure or use sparse matrices. A monolayer object is created using the function create_monolayer_object, which as input can take matrices, edge lists and igraph objects, and can also incorporate node attributes. With a created monolayer object, Infomap is ready to run. A basic example:

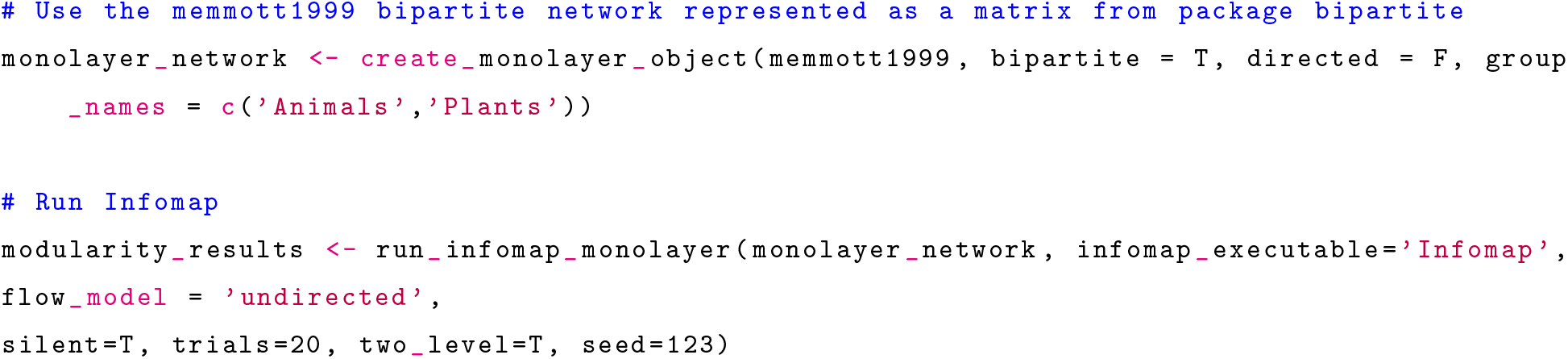

For multilayer networks, the input must be in the form of an edge list. The exact format depends on the existence of interlayer edges. A data frame with nodes is also necessary. It is also possible to provide information on each layer (e.g., coordinates). `infomapecology` will standardise the input and produce a multilayer object with intralayer and interlayer edges, and information on nodes and layers. A multilayer network example:

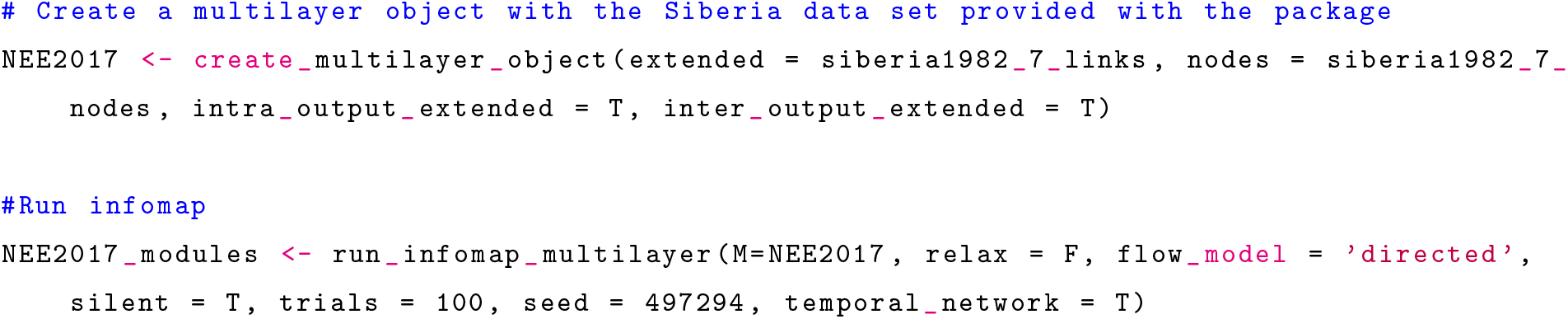

For monolayer and multilayer networks, the results are stored in objects of class infomap _monolayer and infomap_multilayer, respectively, which contain the call for Infomap, the value of L, the number of modules and a data frame with the module affiliation of nodes.

### Use cases

Thanks to its flexibility, Infomap can find modules in many types of networks. Here we exemplify with directed, weighted networks, which are adequate for representing flows, and multilayer networks for analysing modular flows over time. We present other types of networks, including bipartite networks, and hierarchical modularity in the Extended Use Cases (SI 4). The goal of all these use cases is to demonstrate the capacity and flexibility of the framework and to provide general guidelines. We aim to help users analyse their networks rather than to provide full interpretations of the analysed networks.

### Weighted and directed networks

To demonstrate the usefulness of Infomap in identifying flows in weighted networks we use data from Gilarranz et al. (2017), who built an experimental network of 20 cups (patches) connected by tubes and partitioned into 4 modules (Fig. 2). Gilarranz et al. (2017) allowed springtails to disperse freely between the patches and showed that the effects of perturbation to a particular node in the network – leading to local extinction of springtails in the patch – are primarily contained within the cup’s module. Flow modules can provide an adequate description of this dispersal system.

When we applied Infomap and Newman-Girvan’s modularity score Q to the original, unweighted and undirected network (springtails can move in both directions with uniform constraints on movement), both methods partitioned the network into the same four experimentally designed modules. However, when we increased the connectivity between two of the designed modules in the network, Infomap identified 3 modules by merging the two original modules as expected. In comparison, Q still found the same 4 modules (Fig. 2). If we were to repeat the experiment with increased link weights by using wider tubes, we would expect local extinctions to be confined to the 10 nodes within the new module. This ecological example with network flows indicates that Infomap is more sensitive to changes in flows than Q (Table SI 1). (Lancichinetti & Fortunato, 2009; Aldecoa & Marín, 2013) show quantitative comparisons of Infomap and Newman-Girvan’s modularity score optimized with the Louvain method, and (Rosvall *et al.*, 2018) illustrate how the flow-based map equation and the combinatorial modularity score highlight different aspects of networks.

**Figure 2:**
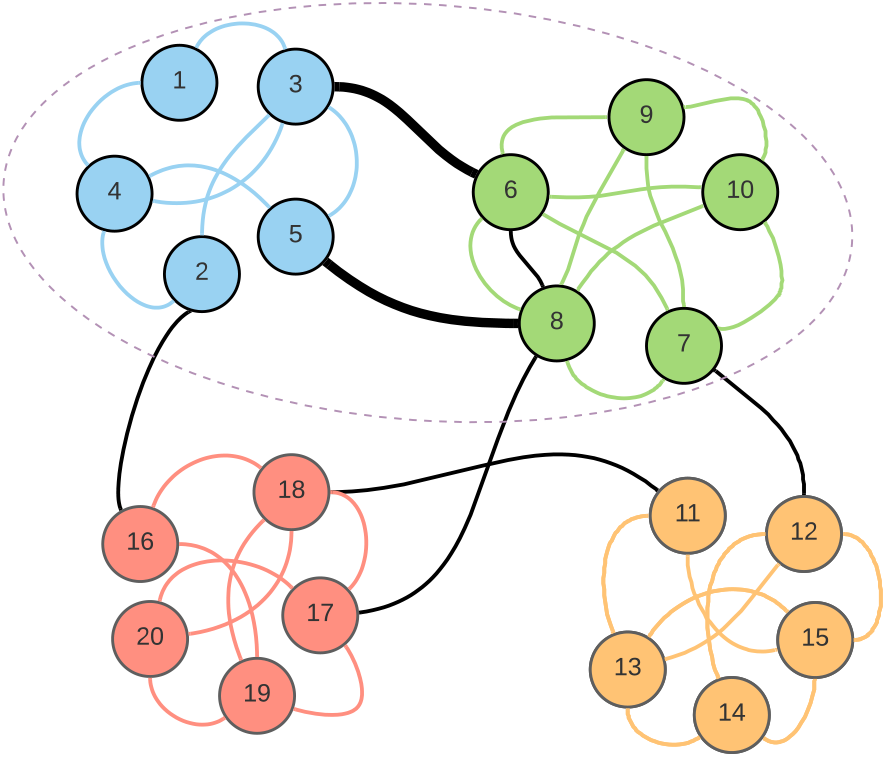
Comparing Infomap to Q in weighted network. The network is the one designed by Gi-larranz *et al.* (2017) to have 4 modules, depicted by node colours. Edges within and between modules are coloured by either module colour or black, respectively and their weight is 1. We computationally increased the weight of two edges between the green and light blue modules (thick black lines) from 1 to 4 in increments of 0.1. This analysis showed that Infomap has a threshold (2.1) above which the two strongly connected blue and green modules merge into a single module (depicted by a dashed ellipse), while *Q* considers them as four modules consistently.

As an example of a directed network, we use data from Tur *et al.* (2016), who measured directed flows of pollen grains (links) in south Andean communities, at three elevations. In their networks, nodes are plant species and links are directed from species *i* to *j* when pollen of species i was detected on stigmas of species *j* (*i* is the donor species and *j* is the receptor). The weights of the links are the number of pollen grains identified. Links between nodes represent pollen movement between species (heterospecific pollination) while self links represent conspecific pollination. Heterospecific pollination occurs when pollinators visit plants of different species and is a cost on reproductive success (see more in Tur *et al.* (2016)). Because the relative flow of self and non-self pollen (con-vs hetero-specific pollination) has ecological and evolutionary consequences, identifying higher-level modules of pollen flow and the roles of particular species in dominating this flow can provide a new perspective into the functioning of this community.

We mapped the pollen movement with and without self-links and found that the structure was considerably different. With self-links, Infomap identified 13 modules, and without self-links 7 (Fig. 3a,b). The increased number of modules with self-loops results from high conspecific pollen flows compared with heterospecific pollination. Because Infomap also quantifies the relative amount of flow at each node, this comparison allows us to look into the roles of individual species. For example, plants that have a large flow of conspecific pollination, but its pollen is also found on many other plants (outgoing flow) are likely to connect between groups of plants through shared pollinators (Fig. 3c).

**Figure 3:**
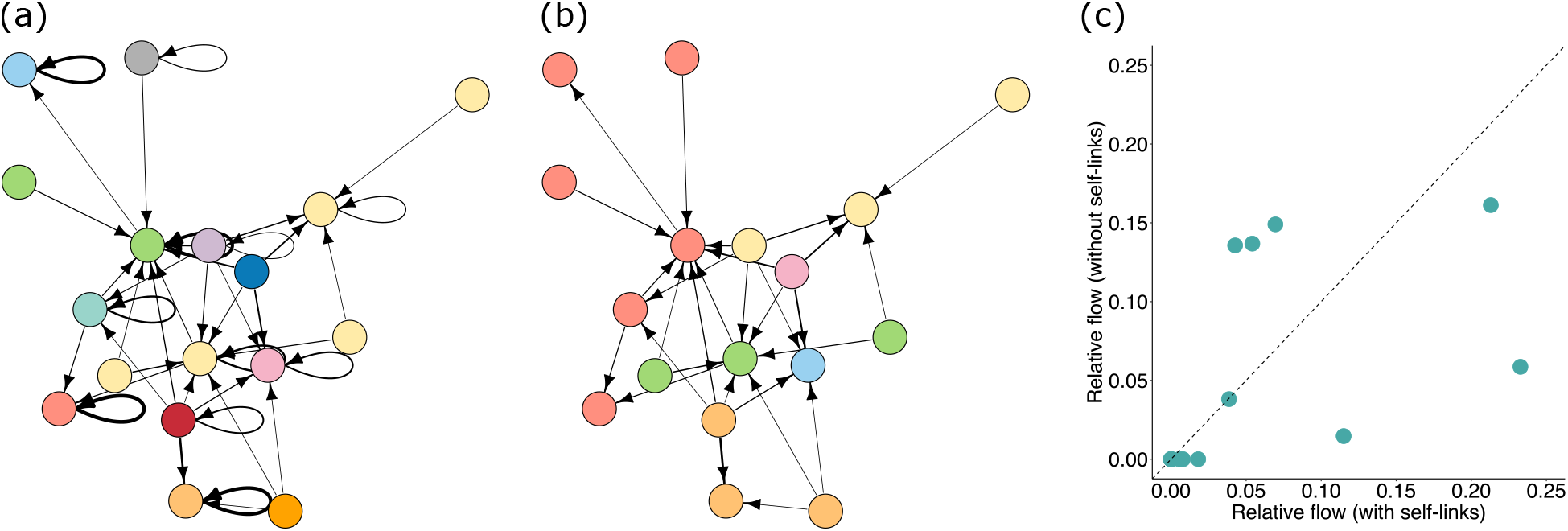
Modularity of a pollen movement network with and without self links. (a) and (b) The network is that of elevation 2000 from Tur et al. (2016). Link width (log-transformed) depicts the mean number of grains of a donor found on a recipient (arrow direction). Node background depicts the affiliation of nodes to modules when self links are not included or included. The position of the nodes in the two networks is the same. It is clear that nodes that are grouped in the same module without self loops are not grouped together in when loops are included. For example, the 6 red nodes in panel b are grouped to 6 different groups in panel a. (c) Comparison of the relative outgoing flow of nodes with and without self-links. The diagonal dashed line represents an equal contribution to flow when self-links are included or not. Plants that are above or below this line have a discrepancy in their contribution to flow, and therefore, structure.

### Temporal multilayer network

There are many types of multilayer networks in ecological systems and the ability of Infomap to integrate layers of different kinds opens up a range of possibilities for their analysis. Per our goal in this paper, we present an example of a temporal multilayer network, which represents flows over time. We use a hostparasite network recorded over six years, in which both interlayer and intralayer links are quantified (Pilosof et al. 2017. The data set is included in infomapecology) and we analyse it in two ways: First, we analysed the network using the existing interlayer links. We found that 47.4% of the modules persisted for all six layers, while 7.89% appeared in only two layers. No module appeared in only a single layer (Fig 4a). This indicates that the grouping of species has a strong temporal component (although we cannot rule out biases due to uneven sampling across time). A second finding is that affiliation of species to modules is flexible: Infomap assigned 21.8% of the species to at least two different modules during the six years. Infomap can assign a species to one module at one time point (layer), and a different module in the next layer because different state nodes represent the same species in different layers (Fig 4b). Biologically, flexibility in module affiliation in this system may capture interannual variation in host and parasite population dynamics.

To illustrate Infomap’s capabilities to model interlayer links, in a second analysis, we ignored the interlayer links and used global relax rates to mimic the typical situation in which interlayer links have not been measured. We limited the relaxation of the random walker between layers to one layer forward, with no backwards relaxation because time has a direction. By systematically changing the value of *r*, we effectively examined the effect of increasing interlayer connectivity on the structure. The higher the relax rate, the more frequent the movement of the random walker between layers, tightening the connection between layers and potentially affecting structure (e.g., creating modules that persist for longer times). While we do detect variation in the number of modules, module composition and persistence, this variation is not considerable (Fig 5). Nevertheless, these results are specific for this network, and we recommend this kind of sensitivity analysis to choose the appropriate relax-rate that best expresses the dynamics of the network. Moreover, the precise definition of interlayer links or the use of relax rates should be one of the primary considerations when analysing multilayer networks (Pilosof et al., 2017; Hutchinson et al., 2018).

**Figure 4:**
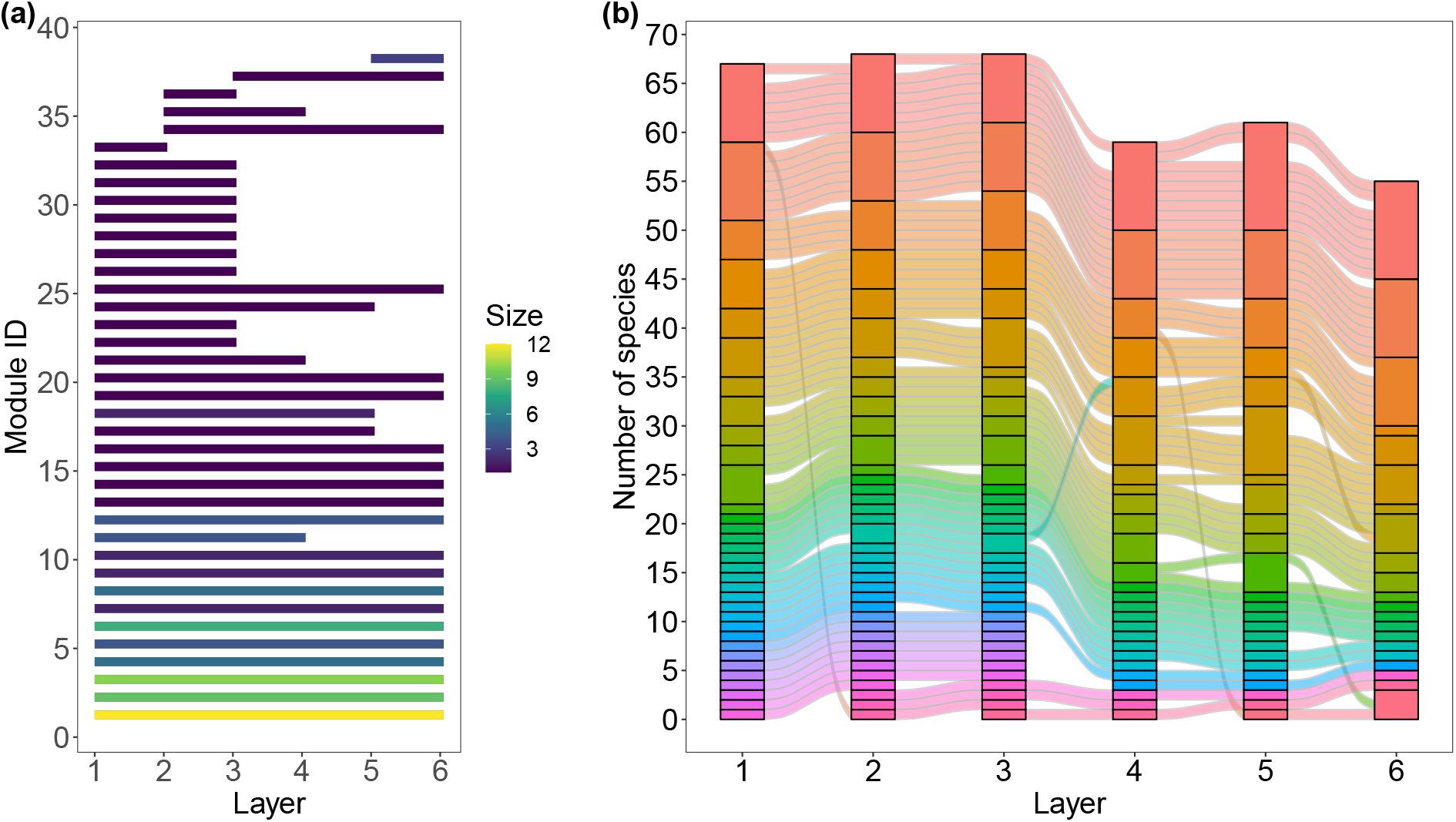
Modular structure of a temporal network. (a) The persistence of each module in time. Colours depict module size: the total number of species in each module. (b) An alluvial diagram for the flow of species from different modules among layers. Species are clustered in modules, presented in coloured blocks. Each line represents a species, and line colours correspond to the module in which the species originates.

**Figure 5:**
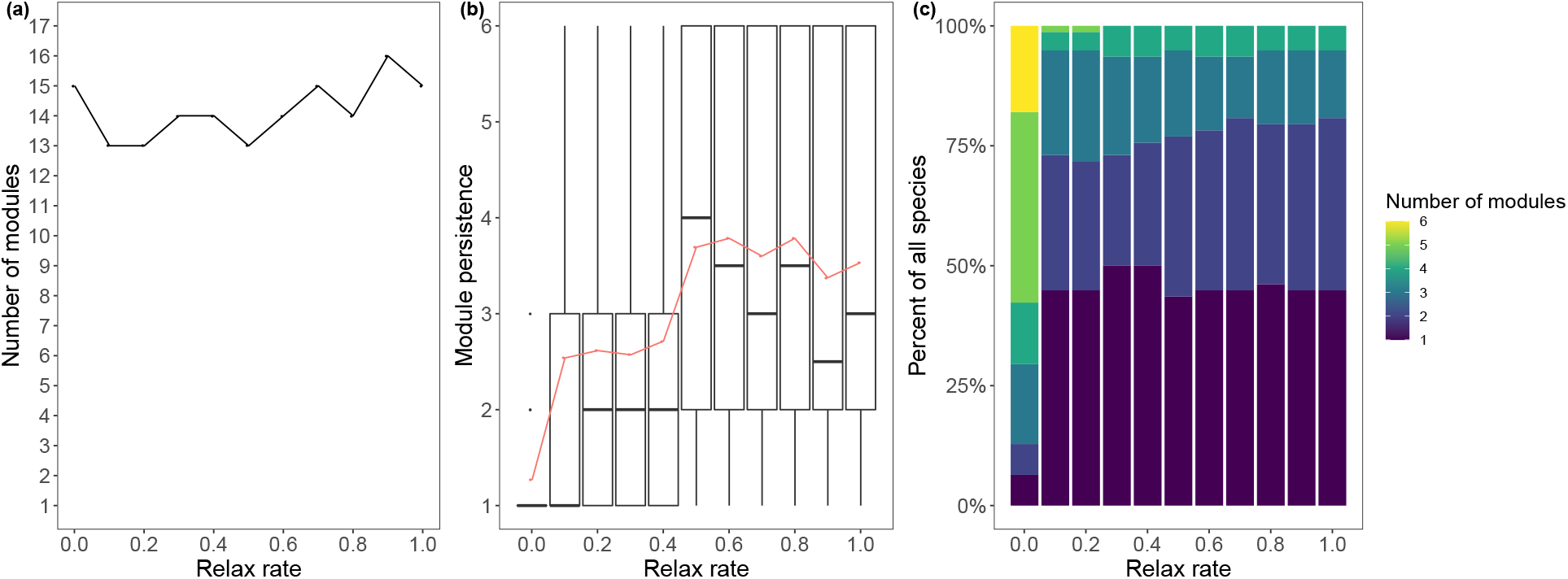
Structure as a function of increasing global relax rate. (a) Variation in the number of modules. (b) Distribution of module persistence (i.e., the number of layers throughout which a module exists). Boxplots represent the range, 95% quantiles and median (black line). The average is marked with red points and line. As expected, module persistence increases with increasing relax rate. (c) Species flexibility: The bars depict the percentage of species appearing in different numbers of modules.

## Conclusion

Modularity is a cornerstone in ecological network analysis because it provides a higher-level simplification of complex ecological systems. Other community detection methods have also shown to be highly relevant for ecological networks, such as stochastic block models which can identify species that are performing unique roles in ecological communities (Sander et al., 2015). Another core concept in research on ecological networks is analyses of the dynamical processes taking place on the network (e.g., Otto et al. (2007)). Nevertheless, the algorithms commonly used in ecology focus on network topology and do not specifically view modules as dynamical building blocks. Here, we aimed to fill this gap by introducing Infomap to ecological research. Modules revealed by different methods (e.g., Infomap or *Q*) will highlight different aspects of networks (Rosvall et al., 2018) (Table SI 1). Infomap, which seeks to coarse-grain the system’s dynamics will identify flow modules, which will likely better capture structural patterns important for the dynamics of the system than other methods.

Like any other method for detecting modules, Infomap can not find a “true” partitioning of a network (Peel *et al.*, 2017) because such partitioning does not exist. We advocate the application of a method appropriate for the question (Table SI 1). For example, if the goal is to detect species that consume, or are consumed by, similar species, then stochastic block models (e.g., the group model (Allesina & Pascual, 2009)) is adequate (Table SI 1). When applied to undirected networks, Infomap provides accurate solutions according to benchmark tests. Nevertheless, Newman-Girvan modularity may be more appropriate if the goal is to detect topological groups by comparing to a random expectation.

The performance and flexibility of Infomap offer several advantages. It is an efficient and fast algorithm, which is particularly useful when analysing a large number of networks (e.g., during hypothesis testing) or large and dense networks. It is also flexible and handles many network types. The possibility of using node attributes to inform the analysis is another advantage (SI 3.5, highly relevant for ecological data, in particular as all interactions rarely are captured in the data (Jordano, 2016). Additional information from other systems, such as information on the role of species traits (Eklöf *et al.*, 2013) and taxonomic classification for interactions (Eklöf *et al.*, 2012), or expert knowledge can then be valuable information for detecting modules.

Modularity has mainly been a theoretical construct in network ecology and empirical work is needed to complement the many generated hypotheses, including the effects on system stability (Grilli *et al.*, 2016; Dormann *et al.*, 2017). As an algorithm specifically designed for coarse-graining the dynamics and identifying flow modules, Infomap is highly relevant for analysing ecological networks (Edler *et al.*, 2017b; Calatayud *et al.*, 2019; Pilosof *et al.*, 2019). The incentives, guidelines and examples presented in this application paper, provide a springboard to take maximum advantage of empirical work in network ecology.

## Acknowledgements

This work was supported by research grant ISF (Israel Science Foundation) 1281/20. D.E. and M.R. were supported by the Swedish Research Council, Grant No. 2016-00796. A.E. was supported by the Swedish Research Council, Grant No. 2016-04919. We thank Ana M. Martín González for advice on data sets and comments on drafts. The authors declare no conflict of interest.

## Authors’ contributions

MR and SP conceived the study; CF, AE and SP collected the data; CF, DE, AE, SP analysed data; CF, MR and SP led the writing of the manuscript. All authors contributed critically to the drafts and gave final approval for publication.

## Supplementary Information

### SI 1 Approaches for modularity analysis

In the following Table SI 1 we provide general guidelines as to which kind of approach for modularity analysis is most adequate in ecological systems. We emphasise that the examples we give are far from exhaustive and that we have not included all of the available tools.

**Table SI:**
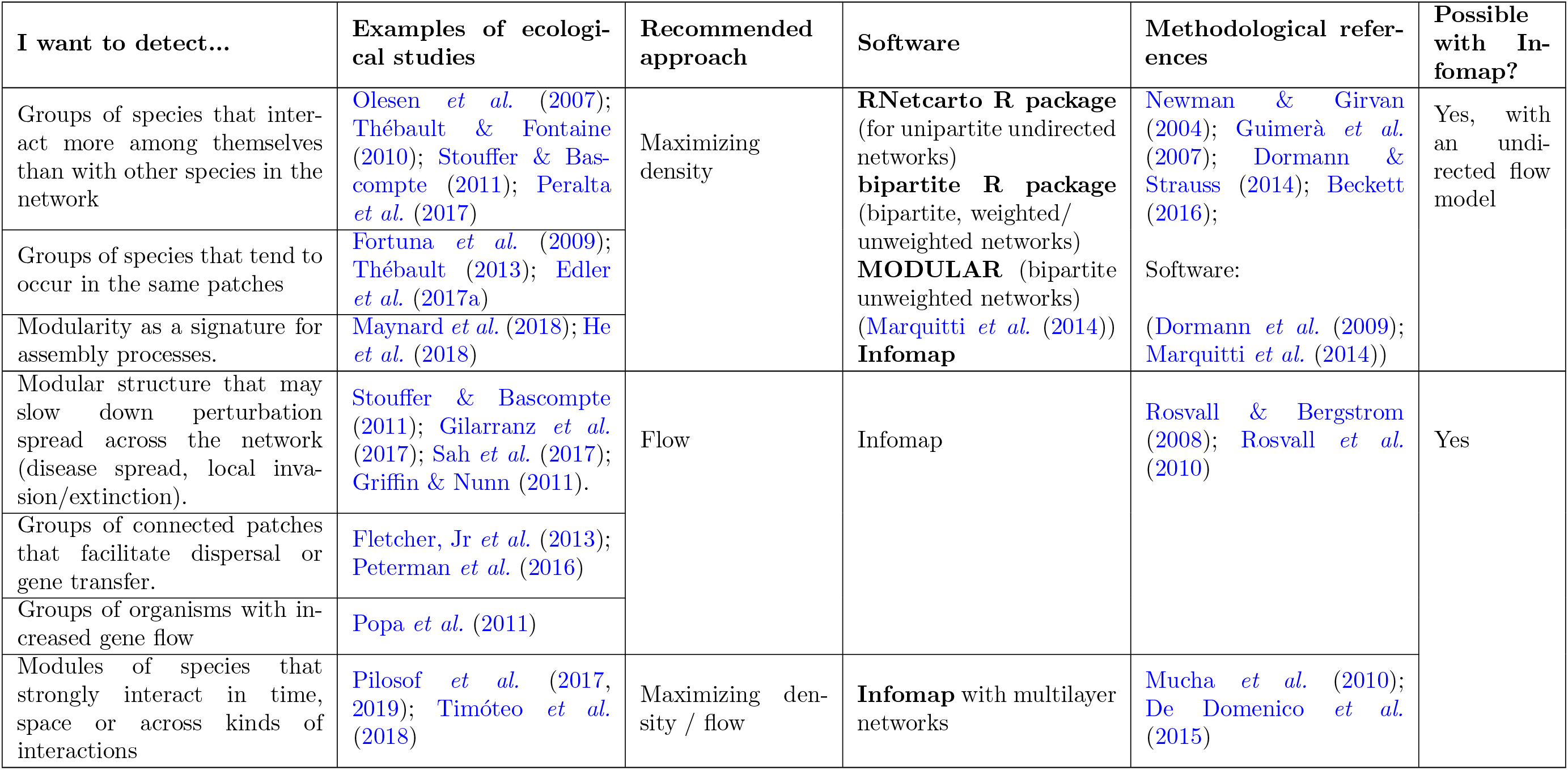
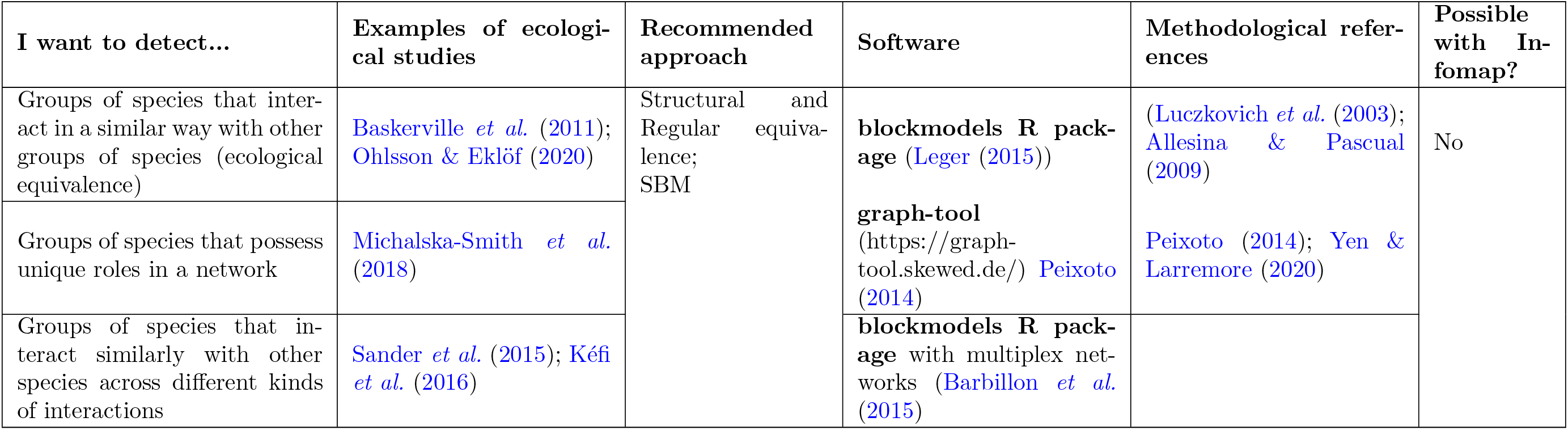
General guidelines for detecting groups (or modules) in ecological systems. We note that: (1) the recommended approach is not necessarily the one commonly applied in the literature; (2) the referenced examples from ecological studies is merely a sample — many more studies exist; (3) there are other aims or research questions that will require modularity analysis and that are not mentioned here; (4) software only includes stand-alone packages (not scripts published for a particular study).

### SI 2 Details on Infomap’s search and optimisation algorithm

For all mentioned types of dynamics and networks, the map equation measures the description length for a given modular partition. To identify the modular partition that reveals most modular regularities and has the shortest description length is a difficult optimisation problem due to the vast number of possible partitions. Like any other network clustering method, it is so difficult that the optimal clustering can be guaranteed only by checking all possible network partitions, which is not feasible even for medium-sized networks. Instead, Infomap is designed to efficiently explore the partition space with respect to the map equation in a greedy and stochastic way (Edler et al., 2017a).

Inspired by the Louvain method for modularity optimisation (Blondel et al., 2008) and similar to the improved Leiden method, Infomap first joins neighboring nodes into modules, subsequently joins them into supermodules, and so on. First, Infomap assigns each node to its own module. Then, in random sequential order, Infomap moves each node to the neighboring module that results in the largest decrease of the map equation. If no move results in a decrease of the map equation, the node stays in its original module. Infomap repeats this procedure, each time in a new random sequential order, until no move generates a decrease of the map equation. Now Infomap rebuilds the network with the modules of the last level forming the nodes at this new level, and, exactly as at the previous level, Infomap joins the new nodes into modules, replacing the previous level. Infomap repeats this rebuilding of the network until the map equation cannot be reduced further. With this core algorithm, Infomap finds a fairly good clustering of the network in a very short time.

Infomap improves the accuracy by breaking the modules of the final state of the core algorithm with submodule movements and single-node movements. In submodule movements, first Infomap treats each cluster as a network on its own and applies the core algorithm to this network. This procedure generates one or more submodules for each module. Then Infomap moves back all submodules to their respective modules of the previous step. At this stage, with the same partition as in the previous step but with each submodule being freely movable between the modules, Infomap re-applies the core algorithm on the submodules.

In single-node movements, Infomap first re-assigns each node to be the sole member of its own module. It then moves all nodes back to their respective modules of the previous step. At this stage, with the same partition as in the previous step but with each single node being freely movable between the modules, Infomap re-applies the core algorithm on the single nodes. In practice, Infomap repeats the two extensions to the core algorithm in sequence, alternating between fine-tuning and coarse-tuning as long as the description length decreases enough.

Finally, because the full algorithm is stochastic and fast, repeated starts from scratch, 100 times or more if possible, increases the chance that the description length of the final partition is close to the global minimum. Moreover, analysing all generated partitions, the so-called solution landscape, can enable more reliable community detection (Calatayud et al., 2019). For example, by introducing a distance threshold between partitions within which a cluster of solutions are considered to be similar, Infomap can restart and generate new solutions – expanding and completing the solution landscaspe – until the probability that a new solution fits within existing clusters is higher than a given accuracy level, say 95 percent.

#### SI 2.1 Hierarchical modules

The algorithm described above will give a two-level solution: one level of modules on top of the level of nodes in the network. However, some networks have a hierarchical modular structure where a multilevel solution would give a shorter description length. Infomap can search for nested modules by running two extra steps. First, it repeatedly applies the two-level algorithm to the top modules to add a super module-level until no coarser super module-level description can be found that decreases the hierarchical map equation. Then, for each top module, it recursively tries to find submodules. For a given module, Infomap’s recursive search clears any existing submodules and finds the coarsest modular description of the sub-network by applying the two-level algorithm followed by the first step of the multilevel algorithm. If this step does not decrease the hierarchical description length, the recursive search does not go further down this branch. Otherwise, it will continue the recursive search for each submodule until no finer modular structure can be found.

### SI 3 The map equation

In this section, we first develop the map equation via a simple example of an undirected network—a detailed version of the example we provide in the main text. We then expand this explanation to different cases such as directed networks in subsequent subsections. For an extensive description of the mechanics of the map equation we refer readers to (Rosvall & Bergstrom, 2008; Rosvall *et al.*, 2010; Bohlin *et al.*, 2014; De Domenico *et al.*, 2013, 2015).

#### SI 3.1 Undirected networks

The map equation uses the principle that any structure in data can be exploited to compress the data (Grünwald & Grunwald, 2007). We illustrate this approach and derive the map equation using a simple example based on Fig. 1. In our example, we consider an undirected network with six nodes and compare a single-module with a two-module solution (Fig. S1).

The map equation takes the flows on nodes and links as inputs to measure the amount of information required to describe random-walker movements on the nodes within and between modules. Hence, we first derive the node and link visit rates. Initially, each node receives a unique identifier called ‘code word’ (see below for details how). *Node visit rates* give the use rate of code words that specify the random walker’s position within modules. *Link visit rates* of the links crossing module boundaries give the use rate of code words that specify transitions between modules. In undirected networks, normalising (dividing) the observed link weights by twice the total link weights gives the link visit rates in each direction (in directed networks there is no need to double the link weights). Summing the incoming flows on each node gives the node visit rates. In the undirected schematic network in Fig. S1a, there are seven links. Assuming that each undirected link has a weight of 1 in each direction, the total incoming link weight is 14. Therefore, each link carries flows of link visit rate 1/14 in each direction. Nodes with two links have a node visit rate of 2/14, and nodes with three links have a node visit rate of 3/14. Based on these rates, the map equation measures the amount of information required to describe flows between and within modules.

While it is possible to compute the modular description length and compression rate without deriving the actual code words given to nodes, for a thorough understanding, we first derive them using Huffman codes (Huffman, 1952). Huffman codes map symbols or events to binary codes with variable lengths such that frequent events receive short codes and infrequent events receive long code words. In this way, Huffman codes can compress event-sequence descriptions. We use the node visit rates to build a Huffman tree that minimises the average code length by repeatedly combining the remaining least infrequent events (Fig. S1c). In a Huffman tree, left and right branches are assigned binary letters of 0 and 1, respectively. The map equation uses these binary letters to assign each node a code word based on the path from the tip of the tree towards the node. Together they form a ‘module codebook’ with unique code words. In the single-module solution in Fig. S1a and c, the unique code words enable an unambiguous description of a walk on the network. For example, the walk C-A-B is encoded by the eight-bit sequence 10000001, 10 for C, 000 for A, and 001 for B. The code words can be concatenated because Huffman codes are prefix-free, that is no code word is the beginning of any other code word. The average code length, *L*^Huffman^ = ∑_*j*_ *P_j_l_j_*, with *l_j_* for the code word length of node *j* weighted by its visit rate *p_j_*, gives the average amount of information in bits required to describe a step on the network (Fig. S1c).

In the two-module solution, we can use the same code words for different nodes in different modules, enabling shorter code words that save information (Fig. S1b and d). However, this description is unambiguous only if we know in which module the walk is taking place. To describe movements within and also between modules, the map equation therefore uses two types of codebooks: one module codebook for each module to describe movements within modules and an ‘index codebook’ to describe movements between modules. For each module *i*, an additional exit code word in the module’s codebook signals that the walker exits the module. In the index codebook, a code word for each module i signals that the walker enters module i. With these two types of codebooks, we can describe uniquely decodable walks.

The lengths of the exit and entry code words depend on the module exit rate and module entry rate, respectively, which are equal for undirected networks. In the two-module solution, the index codebook has code words for “enter green” and “enter orange”, which both occur at rate 1/14 (Fig. S1d). With two modules, the average code length is the sum of all code word lengths in the index and module codebooks weighted by their use rates (Fig. S1d). With the two-module coding scheme, the walk C-A-B is now encoded by the six-bit sequence 010011, 01 for C, 00 for A, and 11 for B. The module-changing walk A-F-D is encoded by 011010111, 01 for C, 10 for exiting the green module, 1 for entering the orange module, 01 for F, and 11 for D. On average, the two-module solution requires 2.57 — 2.43 = 0.14 fewer bits (Fig. S1c and d).

Derived from the node visit, module entry, and module exit rates from the random-walk model, the Huffman coding provides the shortest decodable description of flows in the network. However, to take advantage of the relationship between compression and finding network structure, we are not interested in an actual description but only in estimating the description length. **The map equation allows us to bypass the Huffman encoding to directly calculate the theoretical limit** *L* **of the description length**. From Shannon’s source coding theorem, the per-step average description length of each codebook cannot be shorter than the entropy *H* of its events 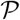, 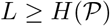 (Shannon, 1948). Using this theorem, the map equation allows us to calculate the average length of the code describing a step of the random walk by weighing the average length of code words from the index codebook and the module codebooks by their rates of use. Summing the terms for the index codebook and the module codebooks, we obtain the map equation, which measures the theoretical minimum description length L in bits given a network partition M (Rosvall & Bergstrom, 2008; Rosvall *et al.*, 2010),

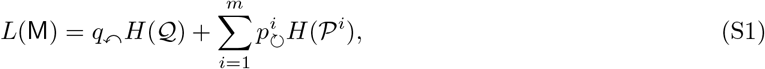

where 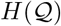 and 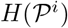 are the frequency-weighted average length of code words in the index codebook and the codebook of module i, respectively. The index codebook is weighted by the rate of entering any module, *q*↶, and each module codebook i is weighted by its within-module flow, 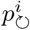, which includes the node-visit rates and the exit rate in module *i*. For the examples in Fig. S1, *L*(*M*_1_) ≈ 2.56 for the one-module solution and *L*(*M*_2_) ≈ 2.32 for the two-module solution. In both cases, L < L^Huffman^, and again the two-module solution requires fewer bits and better represents the network flows.

**Figure S1:**
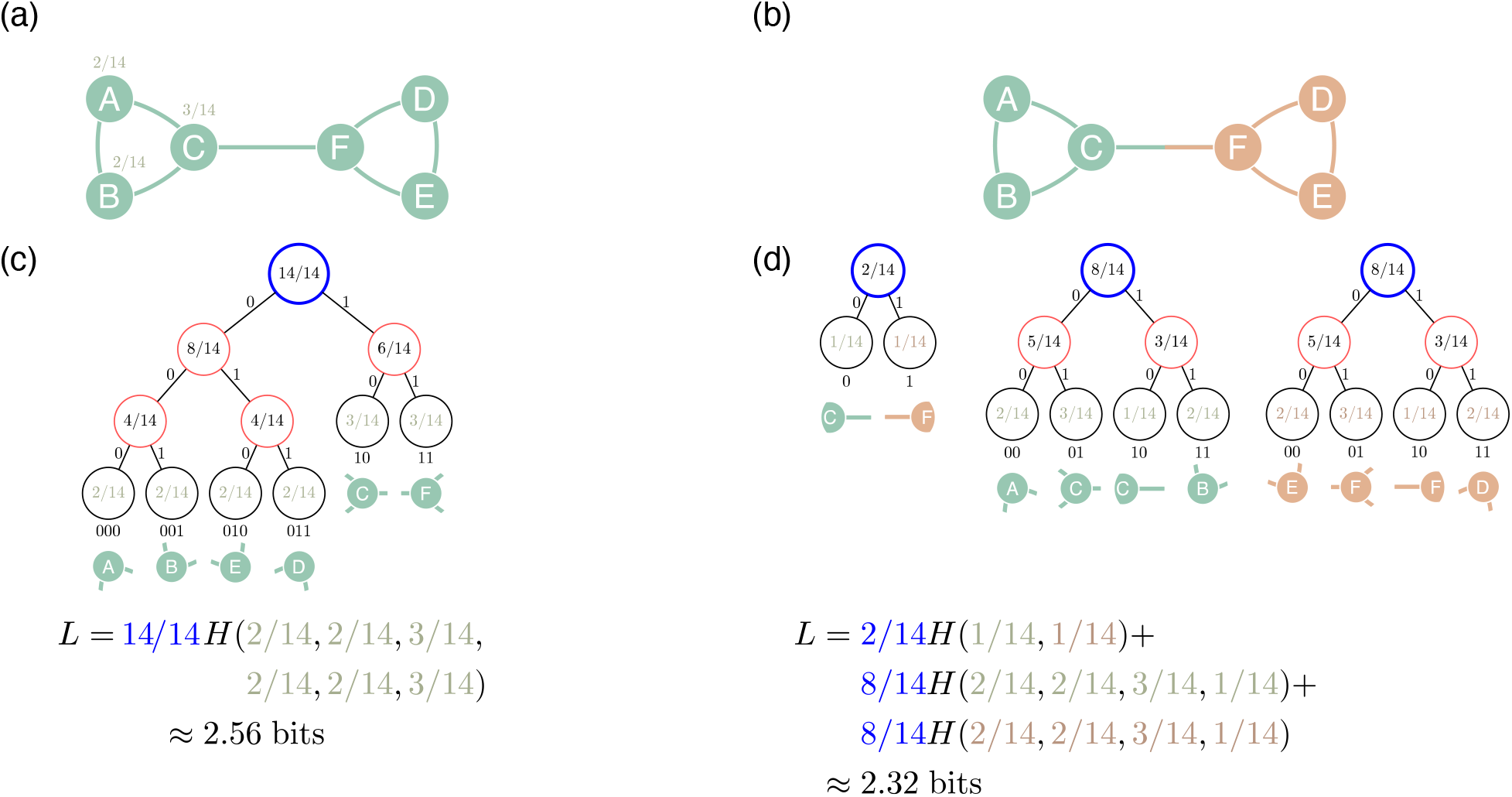
Schematic undirected network. (a) A one-module solution with node visit rates on nodes A-C. (b) A two-module solution. (c) A Huffman tree for the one-module solution. Leaf nodes correspond to nodes in the network. Internal nodes indicate the sum of the node visit rates. Each edge of the tree is assigned a 0 or 1. The code word of each leaf in the Huffman tree is the concatenated 0s and 1s from the top and along the branch of the tree. *L*^Huffman^ is the node-visit weighted average description length required to describe a step on the network using a binary code. (d) The two-module solution uses an index codebook for steps between modules and two module codebooks for steps within modules. In (c) and (d) *L* is the sum of the average code length of each codebook weighted by their use rates and is the theoretical lower limit of the description length.

#### SI 3.2 Directed networks

In directed networks, the procedure for deriving node and link visit rates depends on whether links represent measured real flows or merely constraints on flows. In Infomap, the first option is achieved by setting a flow model as: `-f rawdir`, and the second by: `-f directed`. The available data and research question of interest decide. Suppose we are interested in modelling biomass flow in food webs. Then biomass or energy transfer between taxa measured using the number of prey items in the gut or isotopic signatures are examples of *real flows*. By contrast, binary knowledge on who eats whom merely provides *constraints* on flows. The directed links form interconnected empty channels that direct flows but do not measure flow volumes in themselves. Dispersal rates of an animal between patches is a real flow if we are interested in modelling animal movement. However, if our question concerns the gene flow of that animal, then dispersal rates are constraints on gene flow. Another example is measured contact rates between individuals that can constrain the dispersal of a pathogen.

When link directions and weights represent real directed flows, normalising (dividing) and summing link weights gives link visit, node visit, module entry, and module exit rates in the same way as for undirected networks. However, for directed networks module entry and exit rates need not be the same. For example, assuming that the 20 links in the schematic directed network in Fig. S2 represent real flows, the total outgoing link weight is 20. Therefore, each link carries flows at rate 1/20, nodes with one incoming link have visit rate 1/20, and nodes with two incoming links have visit rate 2/20. For a one-module solution, the entropy of the node visit rates gives the description length 3.92 bits (Fig. S2a). For the optimal four-module solution, the map equation also uses module entry and exit rates. Since both the incoming link to and the outgoing link from a module carry flows at rates 1/20, both the module entry rate and the module exit rate is 1/20 for each module. With one index codebook and four module codebooks, each weighted by their use rate, the description length is 3.10 bits (Fig. S2b). Since the four-module solution requires fewer bits, it provides the better modular representation of the network flows.

When the directed and possibly weighted links represent constraints on flows, we need to model what system-wide network flows the constraints induce. The simplest approach is to derive link and node visit rates from a random-walk model, which requires an iterative process akin to the PageRank algorithm (Brin & Page, 1998). First, each node receives an equal amount of flow volume. Then, iteratively until all node visit rates are stable, each node distributes all its flow volume to its neighbours proportionally to the outgoing link weights. In some cases, however, flows may not be distributed stably. For example, when the flow reaches a node with no outgoing links, it will be stuck in that node, preventing it from reaching others. To solve these issues and guarantee a stable and unique solution, each node teleports some proportion of its flow volume to any nodes proportional to their incoming link weights (see dynamic visualisation available on https://www.mapequation.org/apps/MapDemo.html). This flow redistribution prevents biased solutions.

Because the teleportation steps are not encoded, variations in the teleportation rate have an insignificant effect on the results (Lambiotte & Rosvall, 2012). We note that PageRank is only used for directed networks because it is superfluous for undirected networks. The teleportation rate is adjustable but typically 15 percent. Because the teleportation steps are not encoded, variations in the teleportation rate have an insignificant effect on the results (Lambiotte & Rosvall, 2012). When the nodes’ visit rates are stable, a node’s visit rate times the relative weight of an outgoing link gives the link visit rate. Summing flows on links that exit and enter a module gives the module entry and exit rates as for undirected networks. Because teleportation flows are excluded when deriving link visit rates, module entry and exit rates need not be the same. For a comprehensive description on flow models and of Infomap’s mechanics and how they are applied to different network types we refer readers to Rosvall & Bergstrom (2008); Rosvall et al. (2010); Bohlin et al. (2014); De Domenico et al. (2015).

**Figure S2:**
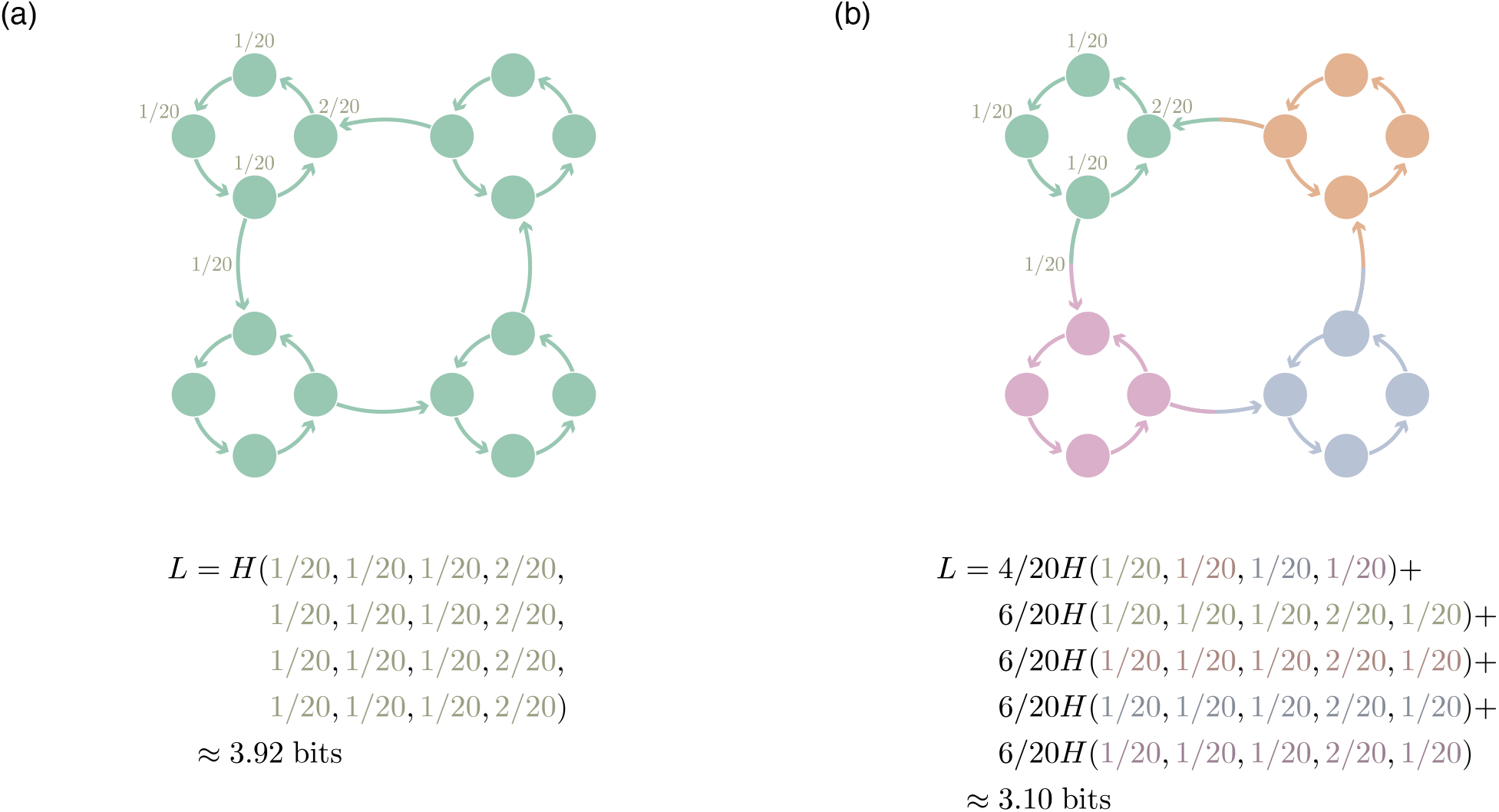
Schematic directed network with real flows. (a) A one-module solution. (b) A four-module solution. The one-module solution has a single module codebook and no index codebook. Therefore, the rates for calculating the entropy are the incoming rates of all the 16 nodes. The four-module solution has one index codebook with four entry rates of 1/20 and four identical module codebooks with three node visit rates of 1/20, one of 2/20, and one exit rate of 1/20. All rates have denominator 20 because each of the 20 links carries an equal amount of flow. The rates of using the module and index codebooks are 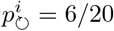 and q↶, = 4/20, respectively.

#### SI 3.3 Bipartite networks

Bipartite networks consist of two types of nodes (e.g. plants and pollinators, hosts and parasites) with links only between the two types. Unlike for *Q*, where specific modification of the objective function to bipartite is required (Guimerà *et al.*, 2007), in Infomap it is possible to consider the network as unipartite, by encoding every step of the random walk (Fig. S3). Nevertheless, there is an extension to this approach (Kheirkhahzadeh *et al.*, 2016). For example, if the research goal is to find modules of plants that tend to co-occur within the same patches, Infomap can encode one type of nodes (plants) by starting the dynamics on that node set and always take two steps between encodings of the walker’s position. This approach will generate modules that only contain plants, though the patches contribute information to that structure. Because in this case Infomap effectively projects the bipartite network into a unipartite network with nodes being separated by two steps between two encodings, the two-step projection typically results in fewer and larger modules compared to encoding every step.

A new development to the map equation does allow however to explicitly consider the bipartite structure by including information on node types (Blöcker & Rosvall, 2020). This requires modifying the map equation and using a different codebook for each node type. In this approach, additional information on node type (worth 1 bit) is added to the map equation. It turns out that even with this additional costly information, this method is better able to compress the description of the network than treating the network a unipartite. The result is better resolution and detection of smaller modules that can potentially reveal deeper network regularities.

**Figure S3:**
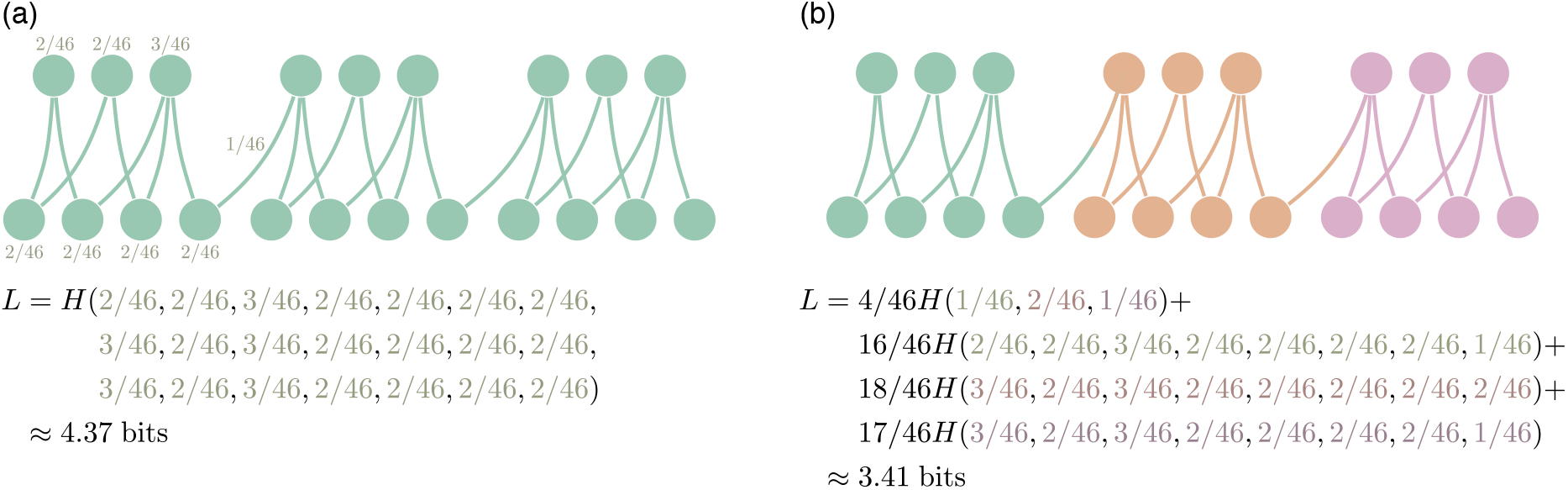
Schematic undirected bipartite network. The description length according to the map equation for (a) a one-module solution and (b) a three-module solution. The bipartite network is treated as unipartite and the modules contain nodes of both types. There are 23 undirected links of weight 1, rendering a total link weight of 46.

#### SI 3.4 Multilayer networks

Multilayer networks enable the representation of multiple layers of data as a single network object (De Domenico et al., 2013; Kivelä et al., 2014). Layers are themselves networks that can represent different points in time or space, or different interaction types such as parasitism and predation. Interlayer links encode ecological processes that operate across layers, such as dispersal, gene flow or population dynamics and represent dependencies between layers. The relative weight between inter- and intralayer links therefore indicates the relative contribution of the processes that operate within and between layers to network structure and dynamics. For example, the relative contribution of movement of individuals within patches and between patches to the structure of dispersal. While the application of multilayer networks to ecology is relatively new (Pilosof et al., 2017), the few studies that investigated multilayer modular structures (not necessarily with Infomap) have revealed new insights into the organisation and dynamics of communities and populations (Pilosof et al., 2017; Timóteo et al., 2018; Pilosof et al., 2019; Mello et al., 2019). Applying Infomap to multilayer networks can capture changes in module evolution and composition over time/space, or capture changing dynamics between different states of the system (De Domenico et al., 2015).

In multilayer networks, nodes representing observable entities such as species are called physical nodes and nodes describing dynamics in the layers for different time points, patches or interaction types, are called state nodes. The random walker moves from state node to state node within and across the layers. However, the encoded position always refers to the physical node (see dynamic visualisation available on https://www.mapequation.org/apps/multilayer-network/index.html). Therefore, the multilayer network representation is not merely an extended network with unique nodes in all layers. Instead, assigning the same code word to all state nodes of the same physical node in the same module acknowledges that only physical nodes represent observable entities and that two state nodes of the same physical node are coupled even if they are not connected. This separation between dynamics and coding provides two advantages. First, it enables a physical node to be assigned to different modules in different layers. From an ecological perspective, this is crucial as, for example, a certain species can have different functions in different layers. For example, in a multilayer network where each layer represents interactions in a different year, a host can interact with different parasites in different years, resulting in temporal variability in parasite spread (Pilosof *et al.*, 2013, 2017). Second, dynamics-coding separation enables to model coupling between layers without interlayer links.

Modelling coupling without provided interlayer links is particularly useful in ecology because interlayer links are often challenging to measure empirically (Hutchinson *et al.*, 2018). If interlayer links are not provided, a random walker moves within a layer and, with a given ‘relax rate’ *r*, jumps to the current physical node in a random layer without recording this movement, such that the constraint to move only within layers can be gradually relaxed (Fig. S4). By tuning the relax rate, it is possible to explore the relative contribution of intra- and interlayer links to the structure.

To relax between layers, Infomap has two options: (1) a rate proportional to the weighted degree of the physical node within the different layers (argument `--multilayer-relax-rate`); or (2) variable coupling based on how similar the connectivity of nodes are between layers (argument `--multilayer-js-relax-rate`) (Aslak *et al.*, 2018). It is also possible to set the number of layers to relax to (argument `--multilayer-relax-limit`). For example, with a relax limit of 2, Infomap can relax from layer 5 to any of layers 3 to 7 (including 5). This flexibility allows Infomap to capture communities in everything from disconnected to fully aggregated layers and from neighbouring to separated layers. Despite the local relaxation, a tightly interconnected group that exists across layers can still form a multilayer community. For example, plants and pollinators that interact consistently across time can form a module that persists for several layers.

Alternatively, Infomap can handle interlayer dynamics in which interlayer links are provided and describe the flow or constraints on flow (using an extended link list input). The system under study, availability of interlayer link data and research question at hand should determine the choice of multilayer dynamics model. This is particularly crucial in case interlayer links are not provided. For example, in a spatial network the natural constraint is free relaxation between layers but in a temporal network relaxation typically occurs towards the next layer because time flows one way.

#### SI 3.5 Considering node attributes

Often ecologists have access to auxiliary information on nodes, which can guide explanation of an observed modular structure based solely on network topology. For example, correlating module composition to phylogenetic distance (Rezende *et al.*, 2009; Poulin *et al.*, 2013) or habitat type (Krause *et al.*, 2003). The opposite approach – to use additional information to guide the community detection – can also be insightful (Newman & Clauset, 2016). This can be particularly important if the interaction network is binary but expert knowledge infer that some interactions are stronger or more relevant. For example, in a seed dispersal network a certain group of species may play a significant role in the actual dispersal of seeds, whereas another group of species interact with the seed (e.g. predation) but do not contribute to successful dispersal (Howe, 2016). A partitioning of the network taking node attributes into account may then give a more accurate description of the true modules of importance for seed dispersal.

**Figure S4:**
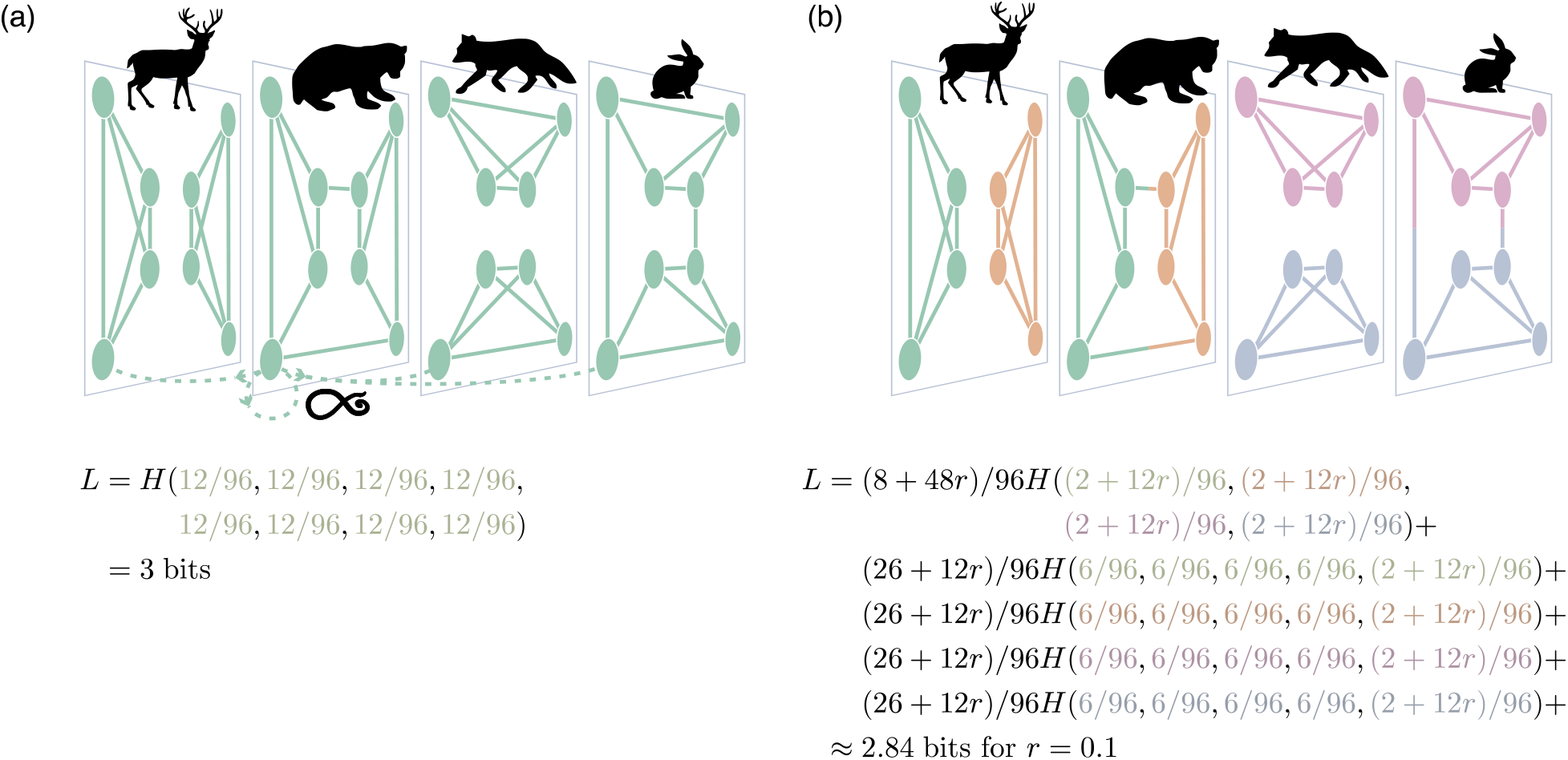
Schematic multilayer network. Within each layer, links represent the dispersal of a species between patches. Each layer is a different species. In the absence of empirical interlayer links, uniform coupling between all layers at relax rate *r* models flows between layers (depicted with dashed arrows for one layer), which is a proxy for the dispersal of a parasite between patches of different species. Note that the random walker can relax to the current layer, as indicated by a self-loop. (a) A one-module solution. (b) The optimal four-module solution. Modules represent patches tightly connected by the parasite’s dispersal on different species.

A recent generalisation of the map equation enables incorporating discrete node attributes in a ‘metadata codebook’ for each module, which accounts for the entropy of the node attributes at each step of the random walk (Emmons & Mucha, 2019). The map equation with metadata (node attributes) takes the form

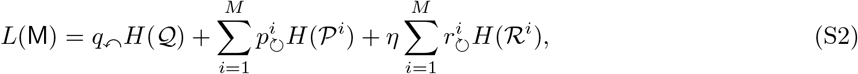

where 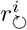 is the total metadata weight of all nodes in module *i* and 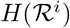 is the module’s metadata codebook. The parameter *η*, adjusted manually, controls the relative contribution of the node attributes to the map equation, with values ranging between 0 for completely ignoring node attributes to infinity for completely ignoring the network structure. Adding node attributes can reveal modules that are not detected by using network flows alone (Emmons & Mucha, 2019).

#### SI 3.6 Hierarchical module structure

Hierarchy is pervasive in all biology, yet hierarchical structures in ecological networks remain little studied. Intuitively, hierarchical modules should reflect structures that are determined by different processes at different scales. For example, in spatial networks, abiotic environmental gradients can determine the top-level modules. In contrast, species interaction such as competition or processes such as local adaptation can determine interactions at lower levels. Alternatively, modules can represent, for instance, a division related to species distribution while submodules represent partitioning according to functional groups (see Olesen *et al.* (2010); Guimarães (2020) for a discussion).

Without limiting the map equation to identify two-level modules, the hierarchical map equation can identify nested multilevel modules that describe hierarchical organisation when such structures enable more compression (Rosvall & Bergstrom, 2011). In the hierarchical map equation, the total code length of every module at all levels gives the total description length. For example, for a three-level map M of *n* nodes partitioned into *M* modules, for which each module *i* has *M_m_* submodules *mn*, the three-level map equation takes the form

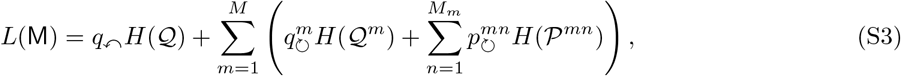

with 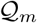 for the exit rate and submodule entry rates in module m at total rate 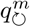, and 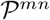 for the exit rate and node visit rates in submodule mn at total rate 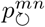. Deeper recursive nesting gives higher hierarchical levels (Fig. S5).

**Figure S5:**
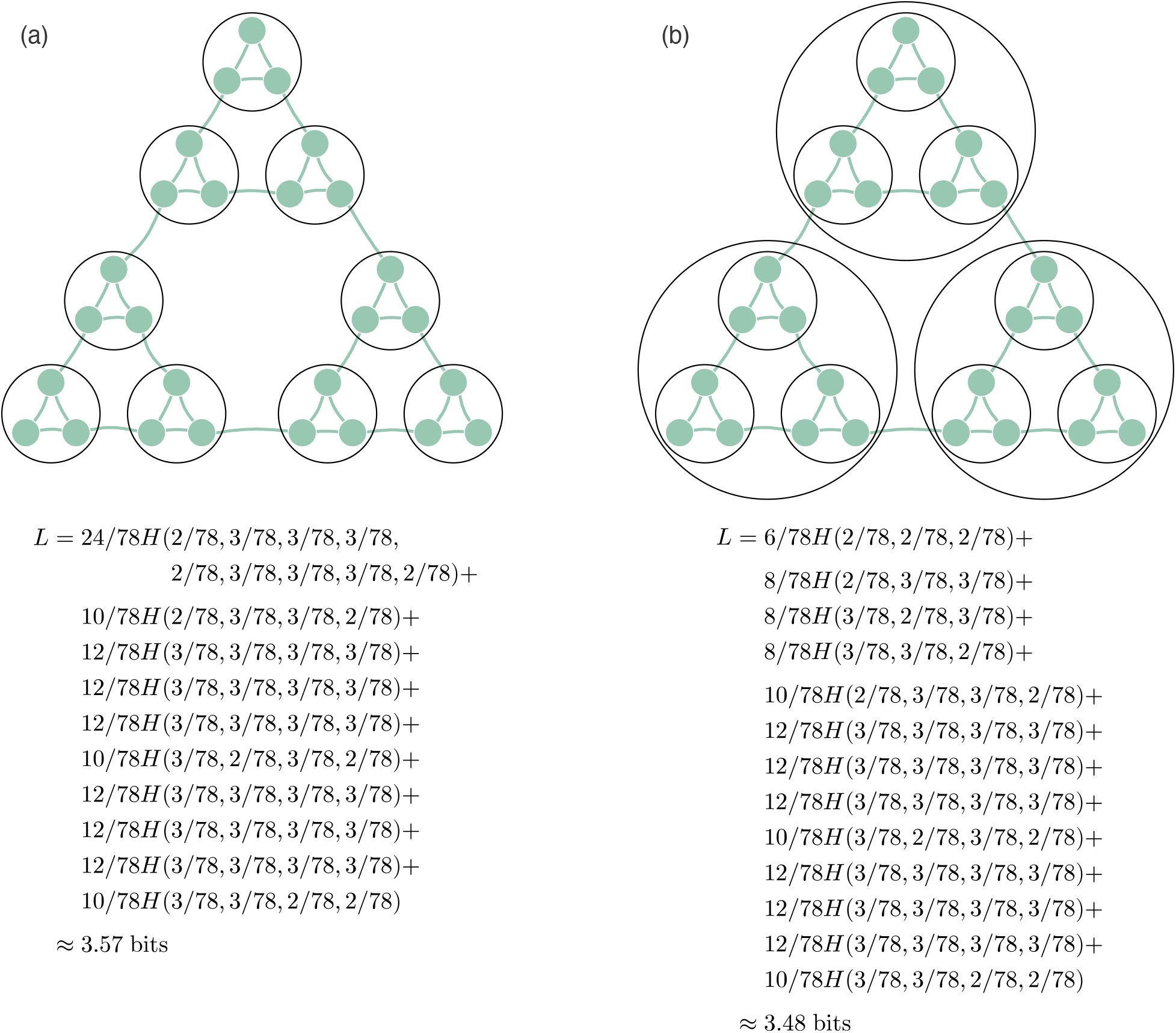
Hierarchical modularity. (a) A one-level solution. (b) A multi-level solution.

### SI 4 Extended use cases

We present use cases for different types of ecological networks to demonstrate the capacity and flexibility of the framework. We focus on the logic behind the analysis of different kinds of networks and present the most relevant input, output and arguments of Infomap. Our aim here, as in the main text, is to guide researchers and not to present full interpretations of the analysed networks, which is beyond the scope of this paper. All technical details such as input/output formats and code are available in https://ecological-complexity-lab.github.io/infomap_ecology_package.

#### SI 4.1 Monolayer bipartite network

We use the bipartite network data set memmott1999 from package bipartite (Dormann *et al.*, 2009; Memmott, 1999). First, we describe the input and output formats of Infomap. The bipartite network is weighted and describes a plant-flower visitor interaction web (25 plant species and 79 flower visitor species) in the vicinity of Bristol, U.K. We use Infomap to detect modules that contain both plants and pollinators. There is evidence that pollinators transmit pathogens between plants (McArt *et al.*, 2014) and that the pollinators share pathogens (Fürst *et al.*, 2014). The number of pollinator visits to flowers - a typical measurement of link weights in plant-pollinator networks - is a constraint for flow of pathogens between flowers (Truitt *et al.*, 2019). Hence, an example of a relevant research question regarding flow would be to detect modules of plants and pollinators between which pathogens are more likely to transmit.

To distinguish between the two node sets, we number the pollinator species from 1-79 and the plants from 80-104. The input to Infomap is described in Table S2. We run Infomap using a flow model for undirected networks (SI 3.3) without hierarchical modules. Infomap’s native output includes module affiliation and the amount of flow that goes through each node as in Table S3 but infomapecology adds the affiliation of modules to the node table (Table S4).

**Table S2:**
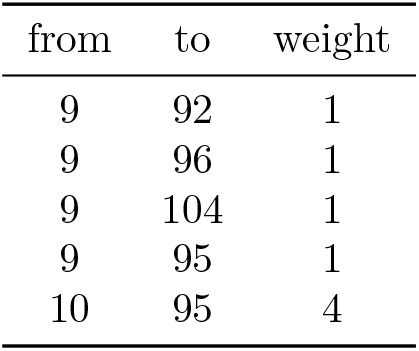
Link list input for a bipartite network. The ‘from’ column is the ID of pollinators (can only contain IDs 1-79), the ‘to’ column is the ID of the plants (can only contain IDs 80-104), and the ‘weight’ column is the frequency visit (link weight).

**Table S3:**
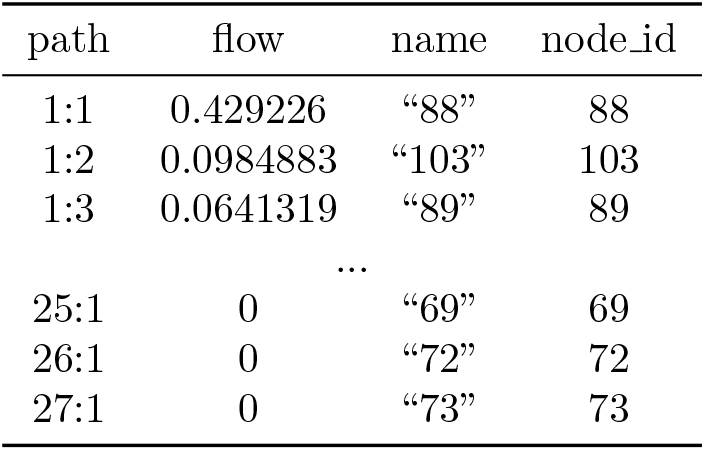
The output of a tree file (first and last three rows). Each row begins with the multilevel module assignments of a node. Colons separate the module assignments from coarse to fine levels, and all modules within each level are sorted by the total flow (PageRank) of the nodes they contain. The integer after the last colon is the ID of the leaf (not the node) within the finest-level module, ordered by flow (in this example, of a two-level analysis, there is only a single colon). The flow column is the amount of flow in that node, its steady-state population of random walkers. The content name column is the node name. The coding scheme in infomapecology incorporates the real node names when parsing Infomap’s output to the final result table (Table S4). Finally, node_id is the index of the node in the original network file.

| path | flow | name | node_id |
| --- | --- | --- | --- |
| 1:1 | 0.429226 | “88” | 88 |
| 1:2 | 0.0984883 | “103” | 103 |
| 1:3 | 0.0641319 | “89” | 89 |
| ... |  |  |  |
| 25:1 | 0 | “69” | 69 |
| 26:1 | 0 | “72” | 72 |
| 27:1 | 0 | “73” | 73 |

**Table S4:** Parsing Infomap’s output. Our code reads the tree output of infomap and integrates it with auxiliary information on nodes provided by the user. Because this is a two-level solution, there is only one level of modules (the second level are the nodes themselves).

| node_id | node_group | node_name | module_level1 |
| --- | --- | --- | --- |
| 1 | A | Aglais.urticae | 1 |
| 2 | A | Apis.mellifera | 2 |
| ... |  |  |  |
| 104 | P | Trifolium.pratense | 2 |

#### SI 4.2 Monolayer directed networks with hierarchical structure

We use the Kongsfjorden food web described in Jacob et al. (2011); Cirtwill & Eklöf (2018a). In this network, nodes are species or taxonomical groups (e.g., Phytoplankton) and binary links represent feeding relationships. Kongsfjorden is a glacial fjord on the northwest corner of the Svalbard archipelago. It is a 30 km open fjord with no marked sill at the entrance, and with a maximum depth exceeding 300 m. The network consists of 262 species with 1,544 feeding interactions.

We allow for hierarchical clustering of modules within modules. We use a directed flow model because the directed links do not represent measured flows of energy. Our analysis shows that the optimal solution has three levels including the leaf nodes (Table S5). To better understand possible mechanisms behind this structure, we use two species characteristics: (1) Feeding type. The species can belong to one of five classes - predator, scavenger, grazer, suspension feeder or none (for primary producers which do not feed on others). (2) Mobility. The species are divided into four classes 1-4 from sessile species to those with the highest mobility. For additional information about the data set, we refer to (Cirtwill & Eklöf, 2018b; Eklöof et al., 2013).

**Table S5:** Infomap output for a three-level solution. The code reads the tree output of Infomap and integrates it with node attributes provided by the user. Here we include three columns with attributes provided in the original data set; FeedingType, Environment and Mobility. The algorithm has partitioned the network into three levels, but the last level is the ID of the leaves within a module. If a module does not have nested modules, it has NA values.

| node_id | node_name | module_level1 | module_level2 | module_level3 | FeedingType | Environment | Mobility |
| --- | --- | --- | --- | --- | --- | --- | --- |
| 54 | Laminaria saccharina | 1 | 1 | 1 | primary | benthic | 1 |
| 55 | Laminaria digitata | 1 | 1 | 2 | primary | benthic | 1 |
| ... |  |  |  |  |  |  |  |
| 31 | Pseudocalanus | 2 | 5 | NA | predator | pelagic | 4 |
| 32 | Oithona similis | 2 | 6 | NA | predator | pelagic | 4 |

Infomap divides the Kongsfjorden food web into four different modules at the first level. The first module contains the majority of species and contains 11 sub-modules at module_level2. The second, third and fourth module do not have submodules in this example. At module level 1 (top modules), the first module is by far the most species-rich and its species’ characteristics are broadly scattered (Fig. S6). A possible explanation for this is that in this network, as in many food webs (Sander et al., 2015), the interactions are mainly between groups of species and not within groups of species (e.g. herbivores eat plants, but not other herbivores). However, if we zoom in to module-level 2, some clear patterns emerge (Fig. S7). For example, looking at the distribution of the species characteristics Mobility and Feeding Type, we see that in submodule 1 most of the species are plants and less mobile grazers and predators (mobility class 2). On the contrary, in submodule 7, the majority of the species are grazers and predators with higher mobility (mobility class 3 and 4). This pattern indicates that the detected submodules associate with different parts of the food web where species interact more closely; highly mobile predators will mainly feed on highly mobile grazers, possibly in spatially different locations from where the less mobile predators feed on the less mobile grazers. An additional example of this differentiation is submodule 8, which contains most of the suspension feeders. Interestingly, most submodules include highly mobile predators, suggesting that this group of species potentially link between submodules (Brose et al., 2005).

**Figure S6:**
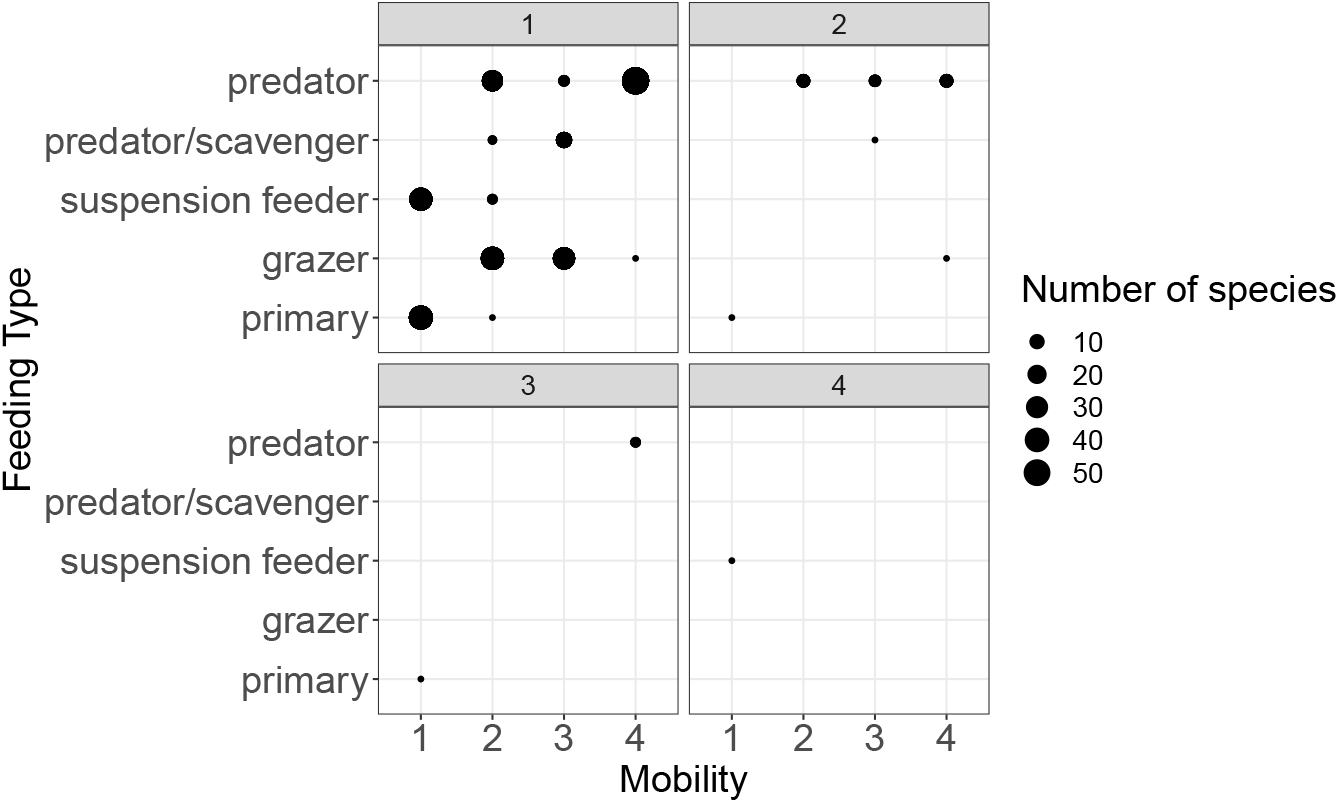
Module composition by species traits for the top modules in Kongsfjorden food web. The x-axis shows the Mobility trait with values between 1 and 4, where 1 means that the species is sessile and 4 is the highest degree of mobility. The y-axis shows the Feeding Type classes. Circle size depicts the number of species in each combination of traits.

**Figure S7:**
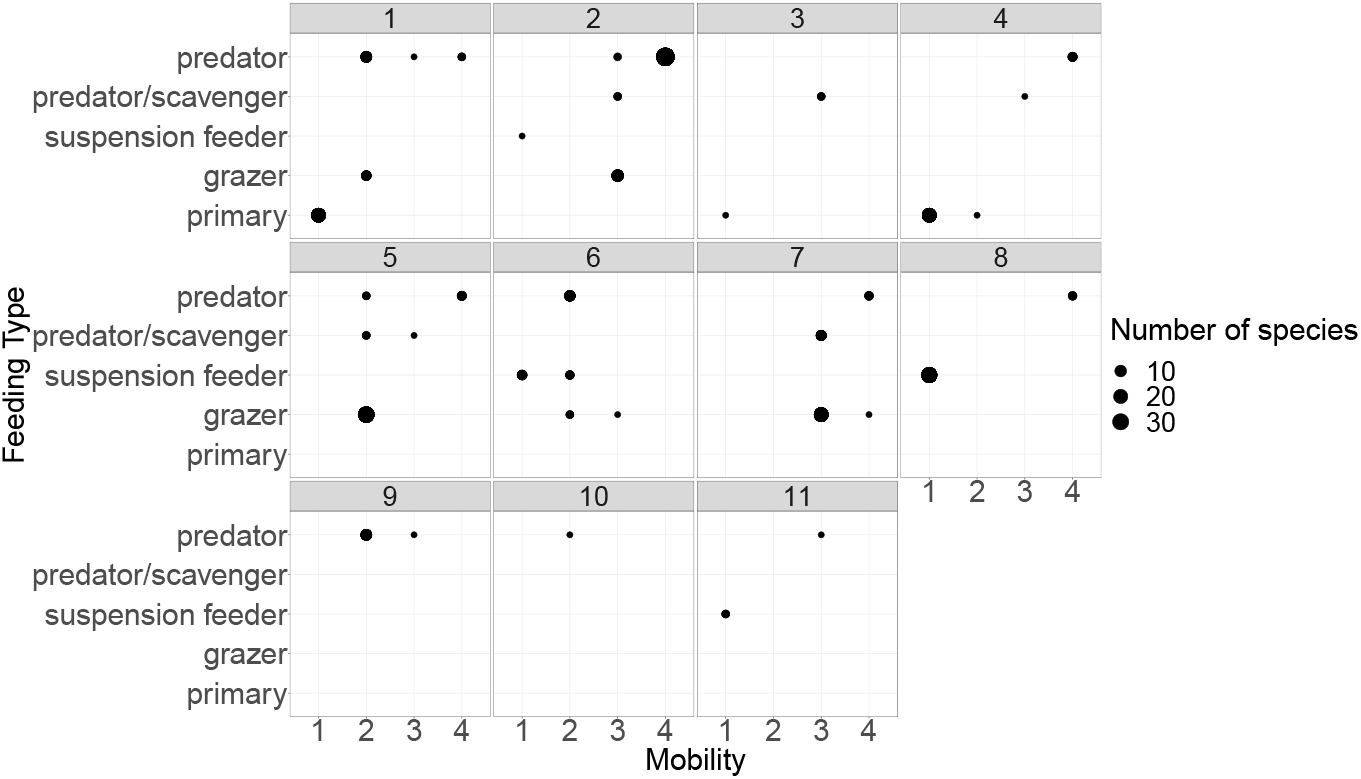
Sub-modules of the first top-module in the Kongsfjorden food web. The figure shows how species with different traits are distributed between submodules within top-module 1. Each panel depicts a different submodule. The x-axis shows the Mobility trait with values between 1 and 4, where 1 means that the species is sessile and 4 is the highest degree of mobility. The y-axis shows the Feeding Type classes. Circle size depicts the number of species in each combination of traits.

#### SI 4.3 Monolayer directed network with node attributes

In this example, we use the Otago food web (Mouritsen *et al.*, 2011). One hypothesis, inspired by the group model of Allesina & Pascual (2009), is that the structure of the network is related to the ‘guild’ of the species because species from the same guild are eaten by the same predators and feed on similar prey. Although the group model identifies both assortative and disassortative groups, the dissortative groups connect to each other and form energy flows in the network (Baskerville *et al.*, 2011) that can be interpreted as groups defined by flows. The organisation of such flow-based groups has been suggested to stabilise the system (Rooney *et al.*, 2008). Mouritsen *et al.* (2011) divided their 140 species to what they term 25 ‘organismal groups’ (analogous to guilds) such as plants, zooplankton, snails, fish or birds. We included these node attributes and tested their effect for different values of *η*, which result in different partitions (Fig. S8). At *η* = 0 node attributes are ignored, and the partition is based solely on network topology. There is a large module that contains 17 of the 25 organismal groups. Increasing *η* returns partitions with more modules, but that contain species from fewer organismal groups. This approach can thus reveal a structure that is *not* supported solely by topology, but rather incorporates expert knowledge.

**Figure S8:**
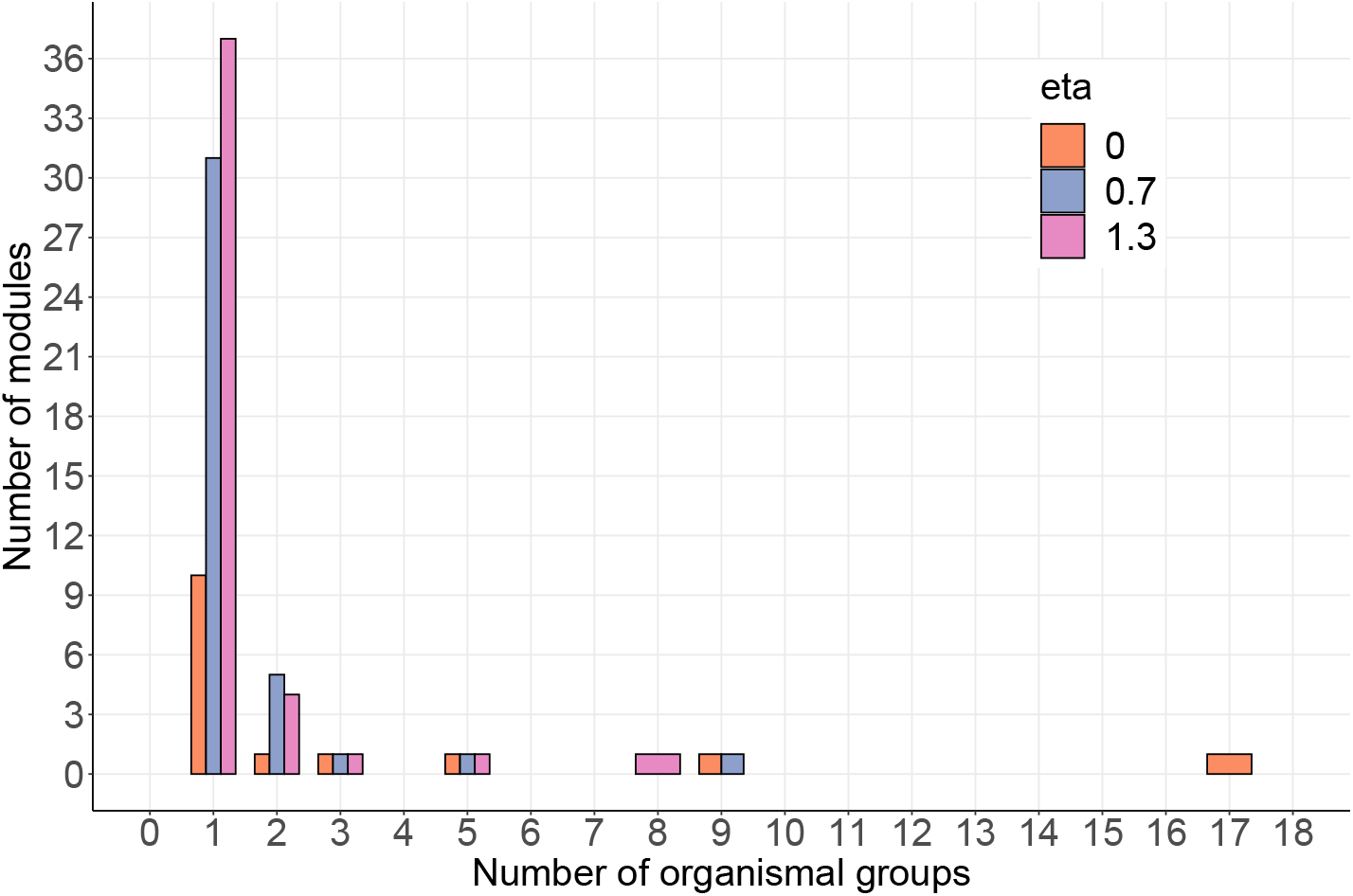
Including node attributes changes network partitioning. The histogram shows the number of modules that contain a particular number of organismal groups (out of 25). With increasing values of η—that is, the more weight is given to the node attributes—more modules contain fewer combinations of organismal groups. For example, there are 10, 31, and 37 modules that contain a single organismal group for η values of 0, 0.7 and 1.3, respectively. Without attributes (η = 0), a single module contains 17 organismal groups. With attributes (η ≠ 0), the maximum number of organismal groups contained in a module is 9.

#### SI 4.4 Controlling flows

As discussed in section SI 3.2, there are two ways to refer to the flow. The first is by measuring real flow; the other is to let instead the links represent some constraints on the flow. Infomap uses real flows with the flow model −f rawdir and constraints on flows with −f directed. A directed flow model will run a PageRank algorithm to set the flows for Infomap, whereas a rawdir flow model will use the flows as specified by the links. We provide a guideline for selecting the correct flow model in Fig. S9.

As an instructive example, we used data from Tur et al. (2016), who measured the directed flow of pollen grains (links) in south Andean communities, at three elevations. In their networks, nodes are plant species and links are directed from species *i* to *j* when pollen of species i was detected on stigmas of species *j* (*i* is the donor species and *j* is the receptor). The weights of the links are the number of pollen grains identified. We focused on pollen movement between species (heterospecific pollination) and did not include self-loops. Heterospecific pollination occurs when pollinators visit plants of different species and is a cost on reproductive success and can affect community structure (see more in Tur et al. (2016)). Flowers between which pollen flows more than to other flowers represent modules.

We selected one network (elevation 2000 m) and ran Infomap twice. First, we used −f rawdir because in this particular example, Tur et al. (2016) state that they “randomly selected five plant individuals per species, whenever possible, and collected five senescent flowers (i.e. post-anthesis) per individual”. Even though this is a sample of the actual population, these are still measured real flows. Hence, a −f rawdir flow model should be preferred. Second, for the sake of example alone, we assumed that the links instead represent constraints and used −f directed. We found that the module assignments of species differ between these two models (Fig. S10). Specifically, using normalised mutual information (Danon *et al.*, 2005), we find that knowledge on the composition of modules in one of the flow models reveals around 77% of the module composition obtained in the other (Fig. S10b). This discrepancy in module assignments indicates that researchers should choose a model carefully, based on the system and sampling scheme. Beyond a careful model selection, comparing flow models may provide insights into the importance of sampling flows. In this example, by sampling only constraints set by pollinator movements, we would miss the existence of a large module (the red module in Fig S10).

**Figure S9:**
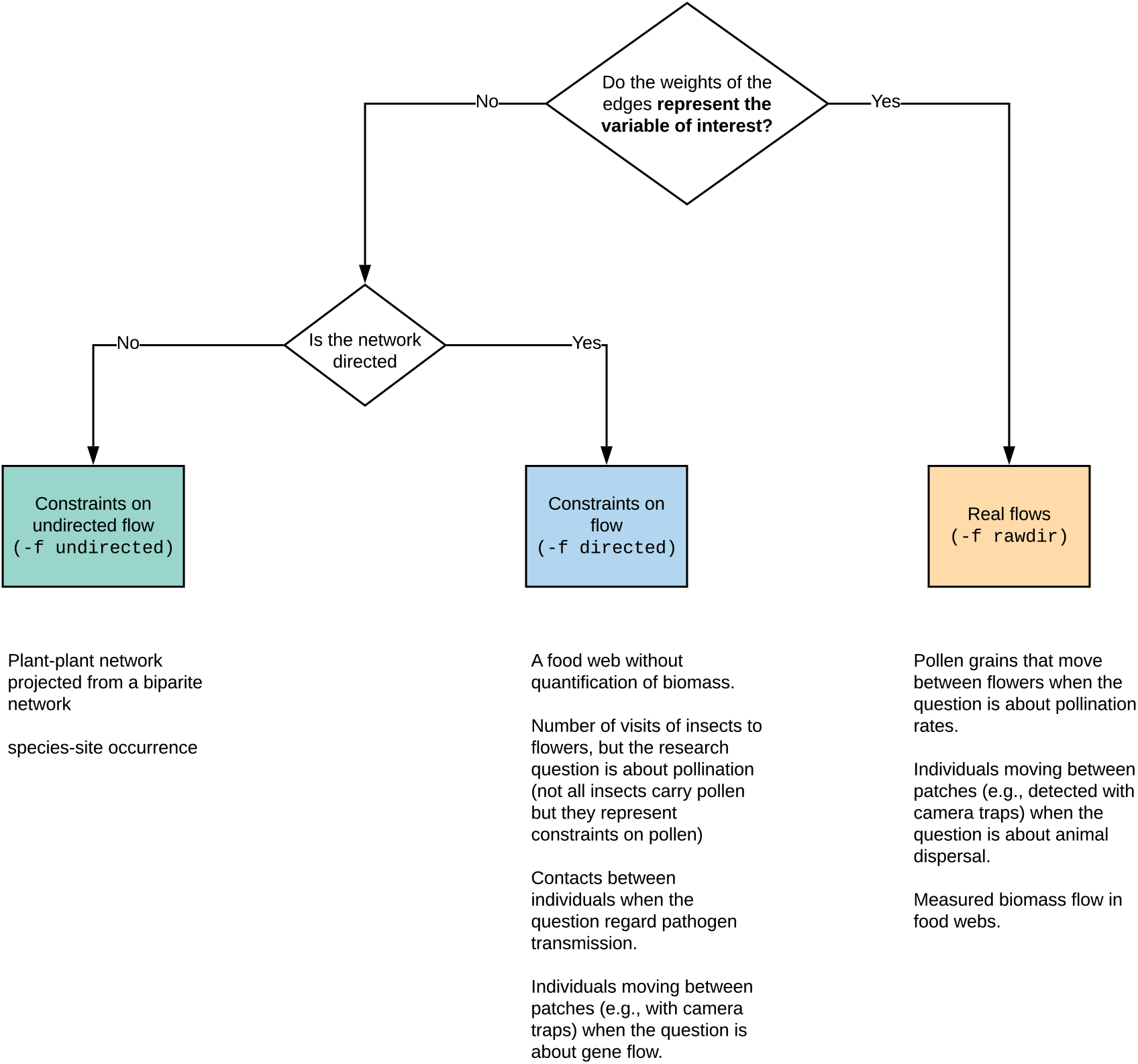
Guidelines for selecting a flow model. The flow model depends on the question and data at hand. We included tangible examples under each flow model.

**Figure S10:**
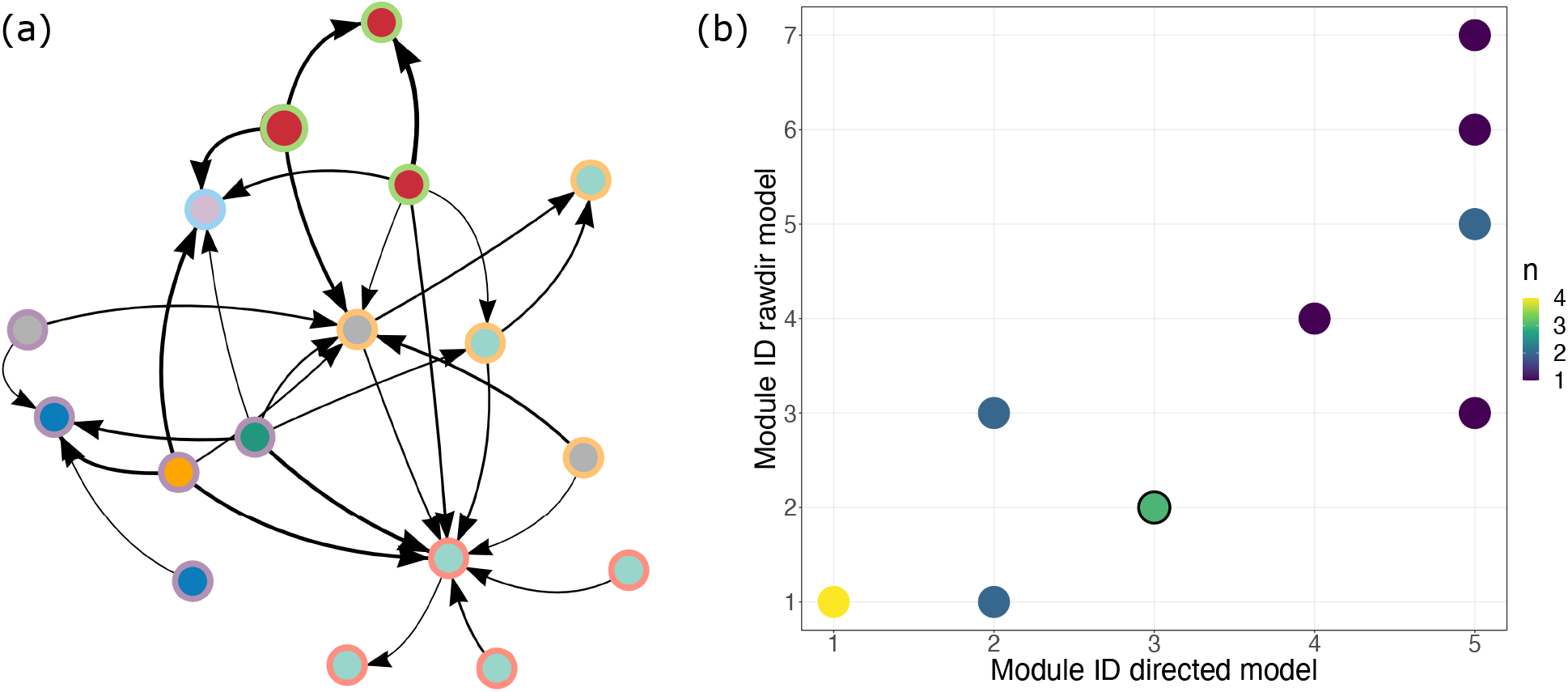
Comparison between flow models. The network of elevation 2000 from Tur et al. (2016)) with no self loops. Link widths (log-transformed) depict the mean number of grains of a donor found on a recipient (arrow direction). Node background and border colours depict the affiliation of nodes to modules according to the rawdir and directed models, respectively. (b) A confusion matrix depicting the number of species assigned to the same module in the two flow models. The colours depict the number of species. For example, the green circle with a black frame depicts three species that were affiliated to modules 2 and 3 in the rawdir and directed flow models, respectively (nodes with a red background and green border in panel a). If module affiliations were identical between the flow models, all circles would lie on the diagonal.

#### SI 4.5 Hypothesis testing

Many objective functions used for community detection, including the map equation, assume that the network data are complete. In practice, however, network data are rarely complete. Consequently, ecologists commonly ask if an observed partition reflects an actual biological process or whether a random network can generate the partition. One approach is to compare the obtained value of the objective function with values obtained by optimising it on a set of randomised versions of the observed network (Gotelli & Graves, 1996; Vázquez & Aizen, 2003; Fortuna et al., 2010). The statistical test measures the probability *P*_value_ of obtaining a network with a better objective function value than that of the observed network. For the map equation, where short code lengths correspond to finding more modular regularities, *P*_value_ = (|*L*_sim_ < *L*_obs_|)/*n*, where *L*_sim_ is the code length of a single randomised network, *L*_obs_ is the code length of the observed network, and *n* is the number of simulations.

The R package infomapecology capitalises on previous methods to shuffle networks from the vegan package, particularly for bipartite networks to automatically run the analysis for hypothesis testing. The user can also use a list of shuffled networks obtained via any other algorithm, as in the example below. Infomapecology also has a dedicated function to plot the results. Here, we demonstrate hypothesis testing using the network of Tur et al. (2016), which we have used in our previous examples in section SI 4.4, for a single elevation (2000 m), with no self-loops. In randomising the network, we explicitly consider the flow dynamics of pollen between plant species. Specifically, we shuffle the network 1,000 times by randomly shuffling the edges between plants. We did so separately for incoming and outgoing links, constraining the total amount of flow that goes into and out of plants as in the observed network but shuffling the plant identities that distribute the flow. This randomisation scheme tests the hypothesis that the way pollen flows in and out between plants is non-random and less modular in simulated networks than observed in nature. We found that the structure of the observed network (Fig. S10) was significantly different than random with a p-value < 0.001 (Fig. S11).

**Figure S11:**
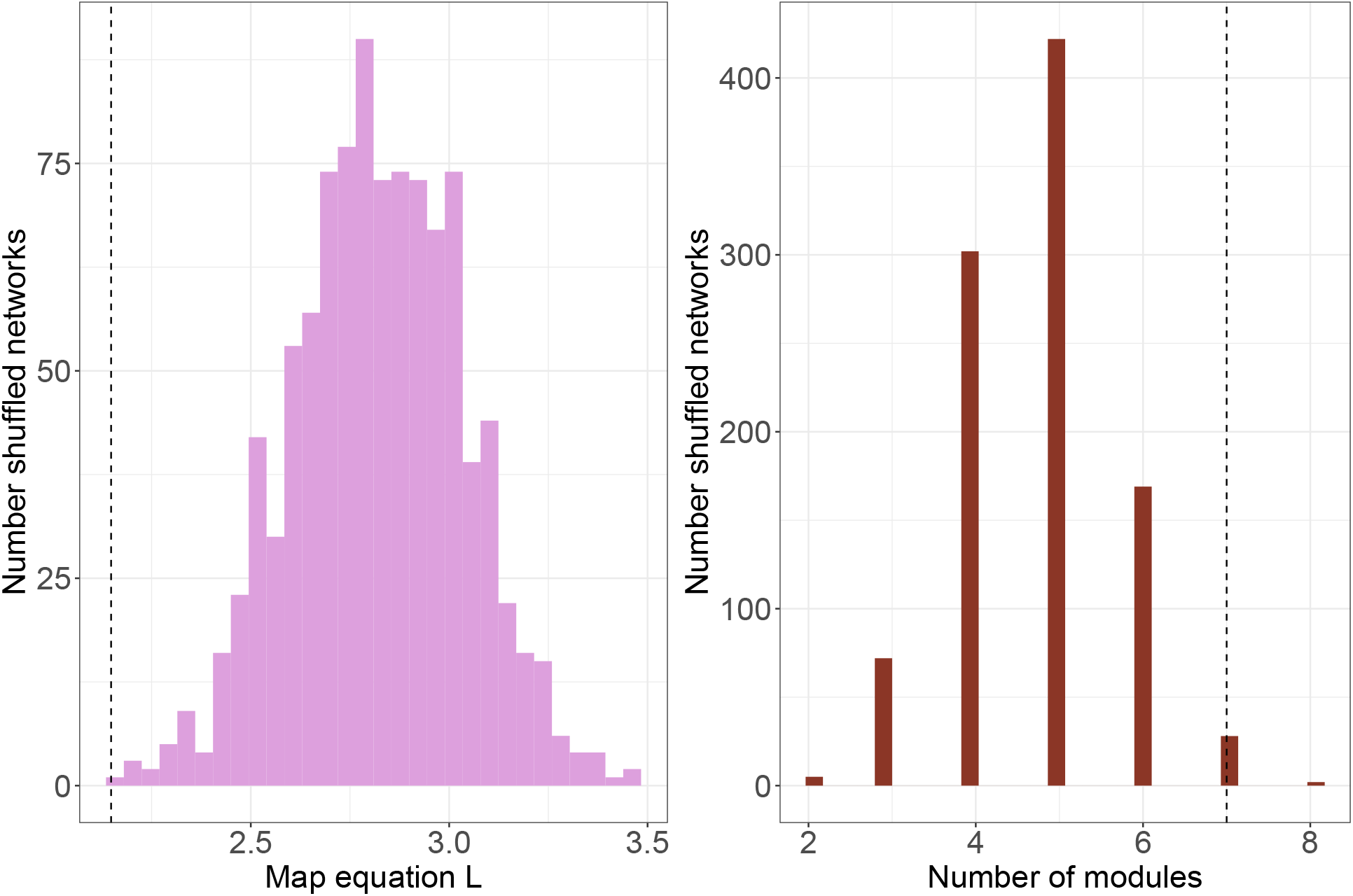
Results of null model hypothesis testing. The plot shows a comparison between shuffled versions and the observed network from Fig. S10. The *L* value of the map equation (a) and the number of modules (b) of the observed network (dashed vertical lines) are statistically significantly different than those obtained for the shuffled networks. The p-values for *L* and the number of modules are *p* < 0.001 and *p* = 0.002, respectively. That is, the network is modular and has more modules than expected at random according to the shuffling scheme.

